# BMAL2 is a druggable target for ARID1A-wildtype ovarian clear cell carcinoma (OCCC)

**DOI:** 10.1101/2025.10.21.683700

**Authors:** Grace Y. T. Tan, Pei-Yi Lin, Li-Tzu Cheng, Yu-Sheng Tsai Yuan, Shih-Han Huang, Chen-Hsin Albert Yu, Chao-Tsen Chen, Peter Chi, Wendy W. Hwang-Verslues

## Abstract

Ovarian clear cell carcinoma (OCCC) is highly chemo-resistant and has worse clinical outcome at advanced stages than other ovarian cancer subtypes. The most frequent (∼50%) alterations in OCCC are AT-rich interactive domain 1A gene (ARID1A) mutations which lead to ARID1A deficiency. However, OCCC that retains ARID1A function differs substantially from ARID1A mutated OCCC. Particularly, targeted therapies that sensitize ARID1A-deficient OCCC to DNA damage are largely ineffective against OCCC with wild-type (wt) ARID1A. Thus, it is important to identify druggable targets and develop targeted therapies specifically for ARID1A-wt OCCC. We identified BMAL2 as a critical OCCC oncogene that promotes tumorigenesis by preventing DNA damage from endogenous origins. BMAL2 depletion altered expression of genes in DNA damage repair pathways, including RAD51, a core enzyme of the homologous recombination (HR) pathway. This led to double-stranded break accumulation, decreased cell viability and reduced tumor growth. This dependence on BMAL2 to maintain DNA integrity and cell viability can be a new route to suppress ARID1A-wt OCCC. Consistent with this idea, we found that GW833972A, a cannabinoid receptor agonist, bound BMAL2 with high affinity and facilitated its degradation. This in turn reduced RAD51 expression, leading to an accumulation of DNA damage and decreased cell viability. Xenograft models further demonstrated that GW833972A treatment alone inhibited ARID1A-wt OCCC tumor growth. Together, our findings reveal an essential oncogenic role of BMAL2 and demonstrate that it is an appealing therapeutic target, especially for ARID1A-wt OCCC.

**Statement of significance:** BMAL2 depletion, or degradation by a small molecule, led to DNA damage accumulation, decreased cell viability and reduced tumorigenesis of ARID1A-wildtype ovarian clear cell carcinoma, indicating that BMAL2 is an appealing therapeutic target for treating this malignant disease.

## Introduction

Ovarian cancer (OC) is highly metastatic and chemoresistant. Due to lack of symptoms and rapid metastasis, most OC cases are diagnosed at late stage (Stage III or IV) when the cancer has spread and the 5-year survival rate is less than 30%. The current standard treatment for OC is complete cytoreduction in combination with platinum-taxane based chemotherapy which can incorporate bevacizumab (1) or Poly (ADP-ribose) polymerase inhibitors (PARPi) (2) as a maintenance therapy after carboplatin and paclitaxel treatment. However, due to complex heterogeneity and chemoresistance, more than 70% of advanced-stage OC patients develop recurrence. Identification of factors that promote these malignant phenotypes is of interest for fundamental OC research and as a means to find new druggable targets.

Five major histological OC subtypes with diverse molecular signatures (3) and therapeutic responses (4) have been classified: high grade serous ovarian carcinoma (HGSOC), low grade serous (LGSOC), mucinous (MC), endometrioid (EC) and ovarian clear cell carcinoma (OCCC) (5–7). HGSOC is the most prevalent subtype, and therefore, has been the most studied. OCCC is the second most prevalent subtype and its incidence rate is rising (8–10); however, our understanding of OCCC tumorigenic mechanisms is limited. While OCCC may be less aggressive in early stages, it has a significantly worse five-year survival (*p* < 0.001) compared to HGSOC. This is primarily due to its de novo chemoresistance in the advanced stage (11). The survival rate for stage III and IV HGSOC patients is about 30%; whereas the survival rate for stage III and IV OCCC patients is ∼30% and 0%, respectively (12). Numerous treatment strategies have been evaluated for OCCC in clinical trials. However, the overall efficacies are not impressive and OCCC subtype-selective effect is low (13,14). Unlike HGSOC, most OCCCs have wild-type (wt) p53 and low BRCA1/BRCA2 mutation frequency (2). The most frequent (∼50%) alterations in OCCC are mutations in AT-rich interactive domain 1A gene (ARID1A) that lead to ARID1A deficiency (2). Treatments that are relatively effective for other OC subtypes, including immune checkpoint blockade, anti-angiogenesis and PARPi are only partially effective, at best, for OCCC (14). Some of these treatments had some effect to sensitize ARID1A mutated (ARID1A-mut) OCCC to DNA-damaging agents; however, the treatments were less effective on ARID1A-wt OCCC (15,16). Thus, identification of druggable targets and development of effective targeted therapies particularly for ARID1A-wt OCCC remains a challenging goal. Detailed characterization of OCCC and how it initiates, progresses and metastasizes will allow more opportunities for OCCC targeted therapeutic development.

Circadian clock genes, including *BMAL1/2*, *CLOCK*, *PER1/2* and *CRY1/2,* form a complex transcription-translation feedback loop (TTFL) to regulate circadian oscillation and contribute to tissue homeostasis in response to the environment (17). These core clock genes are often altered in cancers. Among these clock genes, *BMAL2* (*ARNTL2, MOP9*) is the only one upregulated across different types of cancers (18), suggesting that BMAL2 may have tumor promoting functions. Recently, a few papers reported that BMAL2 expression may facilitate epithelial-mesenchymal transition (EMT) and is associated with invasion and metastasis phenotypes in breast, colon and lung cancers (19–22). However, knowledge about the causal relationship between upregulation of BMAL2 and tumorigenesis is limited.

We identified BMAL2 as a critical oncogene in both ARID1A-wt and ARID1A-mut OCCC. BMAL2 depletion resulted in excessive DNA damage, reduced cell proliferation and viability, and consequently inhibited tumor growth. Mechanistically, loss of BMAL2 significantly altered expression of genes in DNA damage repair (DDR) pathways, including homologous recombination (HR)-mediated repair. One of the BMAL2 downstream targets is a core enzyme in the HR pathway, RAD51. Suppression of BMAL2 reduced RAD51 expression and impaired HR process. Such impact on DNA integrity revealed a new opportunity for ARID1A-wt OCCC patients, as BMAL2 depletion significantly reduced OCCC cell viability. Structural-based virtual screening and additional validations found GW833972A, a small molecule previously identified as a selective CB2 cannabinoid receptor agonist, as a BMAL2 inhibitor. GW833972A facilitated BMAL2 protein degradation, which consequently reduced RAD51 and other DDR genes to increase DNA damage and decrease cell viability. Xenograft models further demonstrated that GW833972A treatment alone can inhibit ARID1A-wt OCCC tumor growth. Together, our findings demonstrated that BMAL2 is a critical oncogene which prevents endogenous DNA damage in OCCC and can be a biomarker as well as a key therapeutic target, particularly for ARID1A-wt OCCC.

## Material and Methods

### Human tissue samples

Eight OCCC formalin-fixed, paraffin-embedded (FFPE) tissue specimens were purchased from Discovery Life Sciences (USA) and a tissue array containing 6 OCCC cases was purchased from Super Bio Chips (CJ3, Seoul, Korea). Sample usage was approved by the Institutional Review Board of Human Subjects Research Ethics Committee of Academia Sinica (AS-IRB-BM-23051).

### Cell lines and culture conditions

OCCC cell lines ES-2 and TOV21G were purchased from Bioresource Collection and Research Center (Taiwan), OVISE and RMG1 from the Japanese Collection of Research Bioresources (Japan), JHOC5 and JHOC9 from the RIKEN Bioresource Research Center (Japan), and OVCA429 was kindly provided by Dr. Noriomi Matsumura (Kindai University, Japan). Normal fibroblast cell line WI38 was kindly provided by Dr. Liuh-Yow Chen (Academia Sinica, Taiwan). All cell lines were authenticated using short tandem repeat profiling (Table S1) and were routinely tested for mycoplasma contamination using a Mycoplasma Detection Kit (BIOTOOLS, TTB-GBC8). OVISE and JHOC9 were maintained in RPMI supplemented with 10% FBS, OVCA429 in DMEM with 10% FBS, JHOC5 in DMEM/F12 with 10% FBS and 0.1mM non-essential amino acids, ES-2 in McCoy5a’s with 10% FBS, TOV21G in 1:1 mixture of MCDC 105 and Medium 199 with 15% FBS, RMG1 in Ham’s F1 with 10% FBS, WI38 in EMEM with 10% FBS. All culture media contained 1x Pen-Strep-Ampho B solution (Biological Industries, 03- 033-1B). Cells were maintained at 37°C in a humidified incubator with 5% CO2 and were used within 25 passages after thawing.

### Reagents

The lentiviral shRNAs, shCtrl (TRC1-Scramble_ASN0004), shBMAL2#1 (TRCN21324) and shBMAL2#2 (TRCN21326), were from the National RNAi Core Facility (Academia Sinica). Cycloheximide (CHX; C1988) and DMSO (D2650) were from Sigma-Aldrich. Olaparib (HY-10162) was from MedChemExpress. RAD51 inhibitor B02 (22133) was from Cayman Chemical.

### Immunohistochemistry (IHC)

FFPE primary tumor and xenograft sections were used for IHC. Sections were dewaxed with xylene and rehydrated with descending ethanol series to water. Antigen retrieval was performed using Target Retrieval Solution, Citrate pH 6.1 (10x), (S1699, Agilent Dako) for 10-25 minutes under high pressure. Endogenous peroxidase was eliminated with 3% H_2_O_2_ for 20 minutes. Slides were blocked with 5% non-fat milk in PBST for 1hr at room temperature, followed by primary antibodies against BMAL2 (PJ90804, Genetex, 1:2,000), RAD51 (GTX100469, Genetex, 1:5,000), γH2AX (9718, Cell signaling technology, 1:500), Ki67 (A20018, Abclonal, 1:12,500), and cleaved-caspase 3 (#9661, Cell signaling technology, 1:200) diluted in blocking buffer overnight at 4°C. After washing, slides were incubated with HRP rabbit polymer for 1 hour before visualization with liquid diaminobenzidine tetrahydrochloride plus substrate DAB chromogen from Dako REAL EnVision (Dako, K5007). All slides were counterstained with hematoxylin. The images were captured under 40X magnification using an Aperio GT450 digital pathology scanner (Leica Biosystems). Positive staining intensity and area were quantified using QuPath digital analysis system.

### Immunoblot (IB) assay

Whole cell lysate was prepared using RIPA lysis buffer (Millipore, 20–188) with 1 mM PMSF (11359061001, Sigma-Aldrich), protease inhibitors (Sigma-Aldrich, S8830) and phosphatase inhibitor cocktail (BIOTOOLS, TAAR-BBI3) followed by sonication at 4°C using a UP50H (Hielscher). Nuclear proteins were extracted using the Subculture Protein Fractionation Kit for Cultured Cells (78840, Thermo Scientific) according to the manufacturer’s instructions to detect γH2AX level. Protein concentration was determined by Bradford assay (5000006, Bio-Rad). Immunoblotting (IB) was performed after SDS-PAGE, with overnight incubation with primary antibodies against BMAL2 (sc-365469, Santa Cruz Biotechnology, 1:500), RAD51 (8875, Cell signaling technology, 1:1000) or γH2AX (2577, Cell signaling technology, 1:1,000) followed by 1:10,000 dilution of horseradish peroxidase-conjugated anti-mouse antibody (115-035-003; Jackson ImmunoResearch, 1:10,000) or anti-rabbit (111-035-003, Jackson ImmunoResearch, 1:10,000). p84 (GTX70220, GeneTex, 1:2000) and GAPDH (60004-1-Ig, Proteintech, 1:10,000) were used as loading controls for nuclear protein and whole cell protein, respectively. Signals were detected using Amersham ECL Select Western Blotting Detection Reagent (RPN2235, Cytiva) and images captured by a UVP ChemStudio Plus BioImaging system (Analytik Jena). The intensity of the bands was quantified using ImageJ software.

### Cellular Thermal Shift Assay (CETSA)

Cells were treated with 20 µM GW833972A or vehicle (DMSO) for 1h at 37°C before subjected to 3-min heat treatment at the indicated temperature (25-60°C). For each temperature condition, 5 x 10^5^ cells in 100uL PBS were used. After heated, the cells were immediately placed on ice for 5 min and then lysed in 0.1% NP-40 lysis buffer (50mM HEPES, pH7.5, 100mM NaCl, 0.1% Igepal, 1mM EDTA) containing protease inhibitors by gentle pipetting. The lysates were centrifuged at 20,000 x *g* for 20 min at 4°C to remove the insoluble aggregates. The soluble fractions were used to determine BMAL2 thermal stability using IB assay after SDS-PAGE. The intensity of the bands was quantified using ImageJ software.

### Drug Affinity Responsive Target Stability (DARTS) Assay

Cell lysates were prepared in 0.1% NP-40 lysis buffer and treated with vehicle (DMSO), 1 or 10 µM GW833972A for 1 h at room temperature with gentle end-to-end rotation, followed by 0.2µg/mL pronase (#10165921001, Merk) treatment for 10 min to induce limited proteolysis. The proteolysis was stopped by addition of SDS sample buffer with 1x protease inhibitor and heated at 100°C for 5 min. BMAL2 protein level was determined by IB assay after SDS-PAGE. The intensity of the bands was quantified using ImageJ software.

### RNA extraction and qRT-PCR

Total RNA was extracted using TRI Reagent (T9424, Merck), treated with DNase I (EN0521, Thermo Fisher Scientific) and cDNA reverse transcribed with Invitrogen SuperScript III Reverse Transcriptase (18-080-044, Invitrogen). qRT-PCR was performed using Applied Biosystems QuantStudio 5 Real-Time PCR System with SYBR GREEN 2X master mix (KM4116, KAPA Biosystems). The mRNA relative quantities were determined using comparative cycle threshold methods with RNA18S5 as an internal control. Primer sequences are listed in Table S2.

### Cell growth, clonogenic and cell viability assays

To determine growth curves, 1×10^5^ cells were seeded in each well of a 6-well plate and cultured under the indicated conditions. Cell numbers were counted at four time points using Olympus R1 Cell counter. Unpaired *t* test was used to test for significant differences between shCtrl and shBMAL2 or vehicle and treatment groups. To evaluate cell survival ability, clonogenic assay was performed using 1×10^3^ cells seeded into 6- or 12-well plates and cultured for at least 7 days. Colonies were fixed and stained with 0.5% (w/v) crystal violet in methanol. Clonogenicity was either measured by colony numbers quantified using VisionWorks software (Analytik Jena), or by extracting the colonies with 15% methanol and 15% acetic acid and measuring the absorbance of the extracted dye at 570 nm. For cell viability assays, 1000 or 2000 cells were seeded in each well of 96-well plates and cultured under the indicated conditions. Cell viability was analyzed using Cell Counting Kit-8 (CCK-8; TEN-CCK8-100, BIOTOOLS) following the manufacturer’s instructions. Absorbance was measured at 450nm using a microplate reader (Sunrise, TECAN). Unpaired *t* test was used to compare shCtrl and shBMAL2 or vehicle and treatment groups.

### Immunofluorescence (IF) staining

Cells were seeded and left to adhere to reach 70-80% confluence on coverslips and then fixed with 4% paraformaldehyde for 20 min at room temperature followed by permeabilization with 0.5% Triton X-100 for 20 min. The cells were then blocked with 1% bovine serum albumin (BSA)/PBS for 1 hour at room temperature and incubated with primary antibodies against phosphor-histone H3 (pHH3; #3377s, Cell signaling technology, 1:800) or γH2AX (#2577s, Cell signaling technology, 1:400) in 1% BSA/PBS for overnight at 4°C. After washing with PBS, the cells were incubated with anti-rabbit CF488A (SAB4600036, Sigma Aldrich, 1:300) secondary antibody for 1 hour at room temperature in the dark. 4’6-diamidino-2-phenylindole (DAPI, D1306, Invitrogen) was used to stain nucleus. Coverslips with stained cells were mounted onto glass slides with fluorescence mounting medium (S3023, Agilent-Dako). Images were acquired using an Andor Dragonfly 202 high speed confocal microscope system and the level of fluorescence signal was measured by fluorescence emission at 505-550 nm with excitation at 488 nm. The fluorescence intensity was quantified using the Imaris 10.1.0 software.

Click-iT EdU Alexa Fluor 488 Imagin Kit (C10337, Invitrogen) was used to evaluate cell proliferation according to the manufacturer’s protocol. Nuclei were counterstained with Hoechst before mounting. Images were acquired using fluorescence microscope (Olympus) and the percentage of EdU-positive cells was quantified using VisionWorks (Analytik Jena).

### Adenovirus-based HR activity reporter assay

Adenovirus-based HR activity reporter assay was performed as previously described (23). Cells were transduced with pAdenoX-pCMV-I-SceI-mCherry alone, pAdenoX-pCMV-DR-GFP-pPGK-ECFP alone or both recombinant adenoviruses, then harvested and analyzed using a LSRFortessa flow cytometer (BD Biosciences) three to four days later. The proportion of EGFP-positive cells was determined to represent HR activity.

### Subcutaneous xenograft model

Animal care and experiments were approved by the Institutional Animal Care and Utilization Committee of Academia Sinica (IACUC# 21-12-1760). Female BALB/cAnN.Cg-Foxn1nu/CrlNarl (NUDE) mice were purchased from the National Laboratory Animal Center (Taipei, Taiwan) at 6-8 weeks of age. Subcutaneous tumor xenograft models were performed with injection of 2×10^6^ ES-2, 5×10^6^ JHOC5 or 5×10^6^ OVISE cells mixed with Matrigel (354234, Corning) with 1:1 ratio into the lower flanks of each mouse. Tumor size was measured every other day using a Peira TM900 tumor volume-measuring device. The experiment was terminated 4 weeks after injection or when tumors reached 2-cm diameter. For drug treatment experiments, mice were randomly assigned to different groups when the subcutaneous tumors reached 50 mm^3^. Tumor growth was monitored every day by Peira TM900. Mice were euthanized when tumors in the control group reached 2-cm diameter or when the tumor-induced skin breakage occurred in the vehicle control group.

### Comet assay

Comet Assay Kit (ab238544, Abcam) was used to evaluate DNA damage. Briefly, cells were embedded in low-melting-point agarose on comet slides and lysed to remove membranes and soluble cellular components. Slides were subjected to alkaline electrophoresis to allow DNA migration. The slides were stained with Vista Green Dye and images were captured using a fluorescence microscope (Olympus). DNA damage was quantified by measuring tail moment using the OpenComet plugin in ImageJ software. At least 50 cells per sample were analyzed in each condition.

### Chromatin immunoprecipitation (ChIP)

ChIP assays were performed using the Zymo-Spin™ ChIP kit (D5210, Zymo Research) following the manufacturer’s instructions. Briefly, cells were crosslinked with 1% formaldehyde for 10 min at room temperature and the crosslinking was stopped by 0.125M glycine. Cross-linked chromatin was sheared using a Bioruptor Pico sonicator (Diagenode) to a size with a range between 100 to 110 bases. Immunoprecipitations were performed with antibodies against BMAL2 (PJ90804, GeneTex) or IgG control (AB-150-C, R&D systems) and ZymoMag Protein A (M2001) magnetic beads. qPCR was performed to detect protein associated promoter regions using primers listed in Table S3. Data were normalized to input and presented as fold enrichment relative to IgG control.

### Structure-based virtual screening of BMAL2 inhibitor

The structure-based virtual screening was performed by TargetMol Chemicals Inc. (USA). The C-terminal PAS2 domain (N360-E477) of BMAL2 resolved by NMR technique [Protein Database (PDB) Entry- 2KDK] and Alphafold protein structure database-predicted BMAL2 full-length 3D structure were used to screen for potential binding pockets. The small molecule binding pocket was detected by using MOE-SiteFinder, where the pocket was served as the docking areas for the virtual screening. Dataset T001, including 18,100 bioactive compounds and natural products, was used to screen potential modulators of BMAL2. The program OEDock (Ver 3.2.0.2) was used for virtual screening, and the compounds with high BMAL2 binding affinity were obtained by the Chemgauss4 scoring function. Based on the docking score, predicted drug-like properties, and binding mode analysis, the top 9 high-affinity compounds with favorable drug properties were selected. Candidate compounds were tested at 5 and 10 μM in cell-based assays to identify BMAL2 inhibitors.

### GW833982A Synthesis

GW833982A was synthesized in-house as outlined in the schematic diagram below. The compound was prepared through a three-step synthetic route starting from the commercially available benzyl 2-chloro-4- (trifluoromethyl)pyrimidine-5-carboxylate (**1**). Reaction of (**1**) with 3-chloroaniline in 1,4-dioxane at room temperature via a nucleophilic aromatic substitution (SNAr) afforded benzyl 2-((3- chlorophenyl)amino)-4-(trifluoromethyl)pyrimidine-5-carboxylate (**2**). Subsequent base-promoted hydrolysis of (**2**) yielded 2-((3-chlorophenyl)amino)-4-(trifluoromethyl)pyrimidine-5-carboxylic acid (**3**), which was then coupled with 4-picolylamine in the presence of EDC·HCl and HOBt·H₂O to furnish GW833982A. All reagents, solvents, and the starting materials were commercially purchased. Benzyl 2- chloro-4 (trifluoromethyl)pyrimidine-5-carboxylate was from Matrix Scientific. 3-Chloroaniline, methylene chloride and 1,4-dioxane were from Thermo Fisher Scientific. 4-picolylamine was from AK Scientific, ethanol (EtOH) was from Shimakyu Chemical, and potassium hydroxide was from Showa Chemical. The reactions were monitored by thin-layer chromatography (TLC) plates using Merck silica gel 60 F254. A handheld UV lamp manufactured by Analytik Jena, UVG-11 (254 nm and 365 nm), was used to identify the spots. All synthesized compounds were characterized by 1H, 13C, and 19F NMR spectra recorded on a Bruker AVⅢ HD 400MHz spectrometer. The chemical shifts are reported in ppm and were internally referenced to the residual solvent signals: for CDCl₃, δ 7.26 ppm for ¹H NMR and δ 77.16 ppm for ¹³C NMR; for DMSO-d₆, δ 2.50 ppm for ¹H NMR and δ 39.52 ppm for ¹³C NMR, respectively. The coupling constants (J) were reported in Hz, and the splitting patterns were singlet,

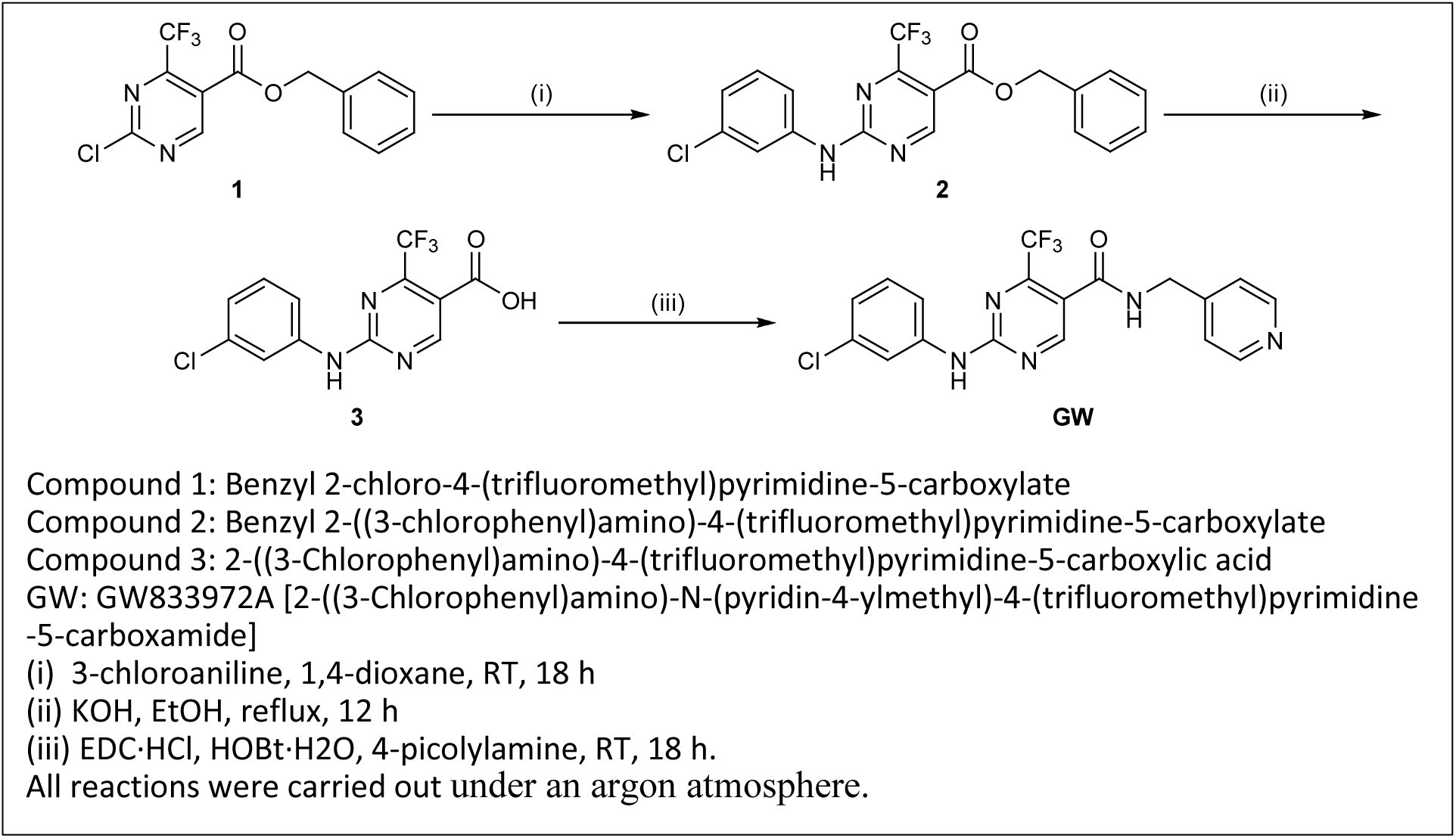

doublet, triplet, quartet, multiplet, and broad peaks, abbreviated as s, d, t, q, m, and bs, respectively. High-resolution mass spectroscopy (HRMS) was obtained on a Bruker microTOF-QII mass spectrometer connected with an Agilent 1260 Infinity HPLC. The ionization source parameters were ESI positive, nebulizer (2.5 Bar), capillary (4.5 kV), dry heater (220 oC), end plate offset (-500 V), scan region (50- 3000 m/z), dry gas (8.0 L/min), and collision cell RF (250.0 Vpp). Detailed structural characterization data can be found in supplementary information.

### RNA sequencing (RNA-seq) analysis

Total RNA was extracted and quantified using NanoDrop (Thermo, ND 1000). All samples with OD 260/280 ratio > 1.8 and OD 230/260 ratio > 2 were sent to BIOTOOLS Company in Taiwan for sequencing. RNA quality was measured by Qsep100 Bio-Fragment Analyzer. Samples with RQN > 6.8 were used for cDNA library construction. Paired-end 150-bp reads, sequenced by NovaSeq 6000 platform, were first processed using Trimmomatic (v0.38) to trim adapters and filter out low-quality bases. The trimmed reads were then aligned to the human reference genome (GRCh38) using HISAT2 (v2.1.0). The Gene-level read counts were subsequently quantified using featureCounts (v2.0.0). After normalization of the count data, differentially expressed genes (DEGs) between shBMAL2 and shCtrl for each cell line were determined based on an adjusted p-value < 0.05 and an absolute log2 fold change > 1 using the R package ’DESeq2’ (v1.26.0). To elucidate the biological pathways associated with the identified DEGs, Kyoto Encyclopedia of Genes and Genomes (KEGG) enrichment analysis was conducted using the R package ’clusterProfiler’ (v3.14.3). For each cell line, the expression profile of each shBMAL2 clone was compared with the shCtrl and the expression fold-change of genes in the KEGG-HR pathway (hsa03440) was calculated (Table S4).

### Statistics

All data are presented as means ± SD (n = 3 unless otherwise noted). *t* test, two-way ANOVA or nonlinear regression (curve fit) analysis was used to compare control and treatment groups. Kaplan–Meier estimation method was used for overall survival and progression free survival analyses, and a log-rank test was used to compare differences. Statistical analyses were performed using Prism 10.

### Data Availability

The RNA-seq data generated in this study are publicly available in Gene Expression Omnibus (GEO) at GSEXXX. (Due to government shutdown, submission high-throughput sequence data to GEO is currently not processed.)

## Results

### BMAL2 upregulation is observed in OC, including the OCCC subtype

*BMAL2* expression is extremely low in normal ovarian tissues (Fig. 1A). However, close to 50% of OC (TCGA-OV) cases in the Cancer Genomics Atlas (TCGA) database have increased copy number of *BMAL2* (Fig. 1B). This is substantially higher than other circadian clock genes and suggests that these tumors have increased BMAL2 expression. Kaplan-Meier (K-M) analysis (24) further showed that OC patients with high *BMAL2* expression tumors have significantly worse overall survival (OS) and reduced progression free survival (PFS) compared to patients with low *BMAL2* expression tumors (Fig. 1C, 1D), indicating a positive correlation between BMAL2 expression and OC malignancy. To determine whether BMAL2 is differentially expressed in OC histological subtypes, the CSIOVDB microarray gene expression database was used and the results showed that OCCC cases have significantly higher *BMAL2* expression compared to SOC (HGSOC and LGSOC) and EC subtypes (Fig. 1E). We then performed immunohistochemistry (IHC) using 14 OCCC primary specimens and found that BMAL2 was exclusively expressed in OCCC cells, not in the adjacent non-tumor cells (NT) (Fig. 1F), indicating that BMAL2 has potential prognostic value and oncogenic roles in OCCC.

**Figure 1.**
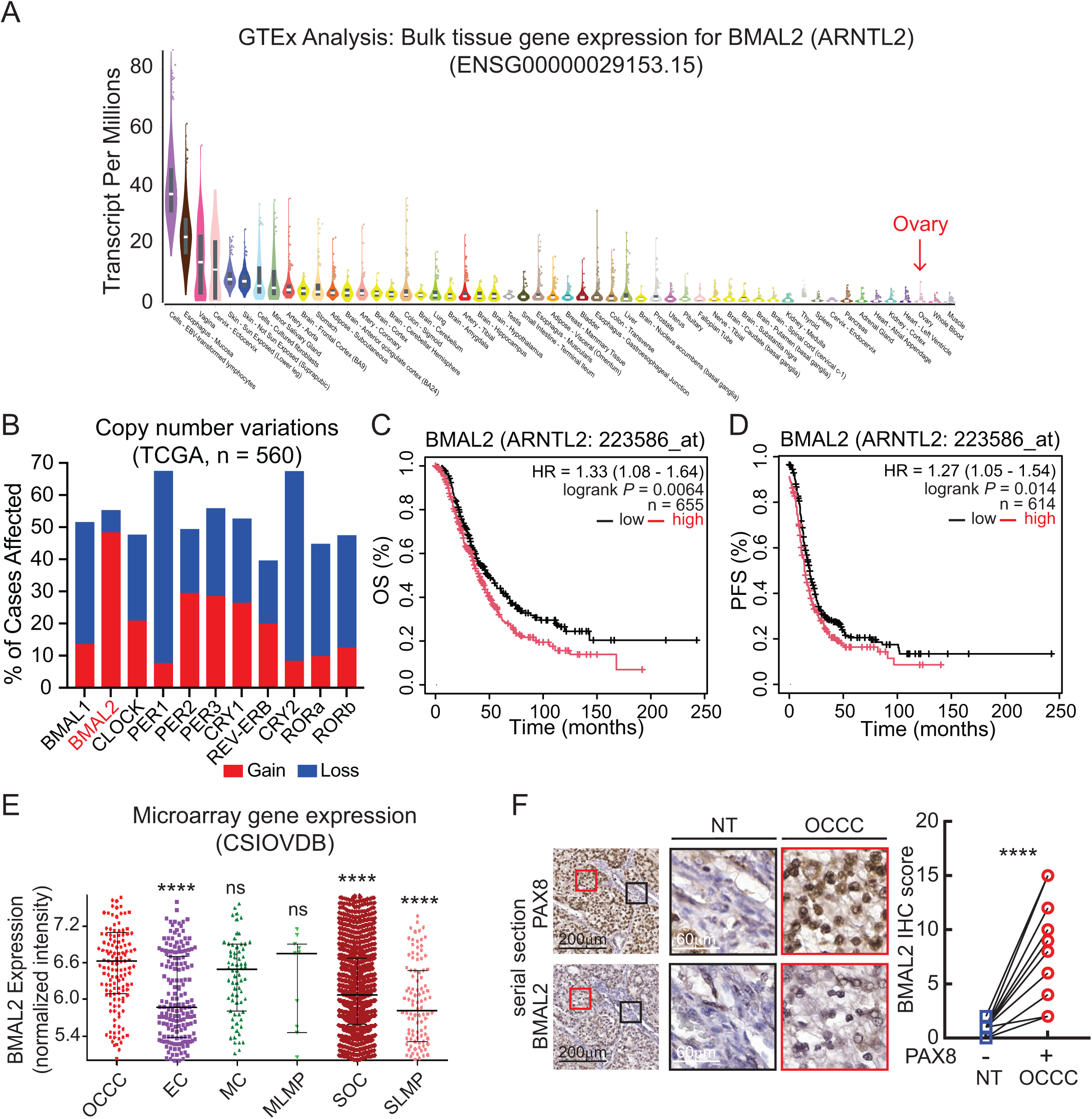
High BMAL2 expression is observed in OC, including OCCC, and correlated with poor clinical outcome. **A.** BMAL2 (ARNTL2) gene expression in normal tissues. The data were obtained from the GTEx Portal (dbGaP Accession phs000424.v8.p2). **B.** Copy number variation of BMAL2 in OC (TCGA-OV) cases. **C.** Kaplan−Meier overall survival (OS) analysis of OC patients grouped by BMAL2 expression. BMAL2- high group (n = 372) is indicated by red line; BMAL2-low group (n = 283) is indicated by black line. *P* = 0.0064. The *P* value was determined by log-rank test. **D.** Kaplan−Meier progression free survival (PFS) analysis of OC patients grouped by BMAL2 expression. BMAL2-high group (n = 215) is indicated by red line; BMAL2-low group (n = 399) is indicated by black line. *P* = 0.014. The *P* value was determined by log-rank test. **E.** BMAL2 gene expression in OC histological subtypes analyzed using CSIOVDB database. The *p* value was determined by Mann Whitney test. ****, *P* < 0.0001; ns, not significant. OCCC, clear cell carcinoma; EC, endometrioid carcinoma; MC, mucinous carcinoma; SOC, serous carcinoma, LMP, low malignant potential. **F.** (Left panel) Representative images of PAX8 and BMAL2 IHC staining using serial tumor sections from OCCC patients. Red boxes indicate enlarged tumor regions. Black boxes indicate enlarged non- tumor regions. (Scale bars, 200 μm or 60 μm as shown in the images.) (Right panel) Quantification and comparison of BMAL2 level in non-tumor (NT) and OCCC tumor regions of 14 OCCC patient samples. Significant difference is based on paired t-test. ****, *P* < 0.0001.

### BMAL2 depletion inhibits tumorigenic ability in OCCC cells

Approximately half of OCCC cases harbor mutated ARID1A while the other half expresses wt ARID1A (2). Since ARID1A is an accessory subunit of the SWI/SNF chromatin remodeling complex, transcriptomic profiles between these two subsets of OCCC differ from each other, and, consequently, the mechanisms driving tumorigenic ability and drug response can also be different. We examined a panel of OCCC cell lines and found that BMAL2 expressed in most of the OCCC cells regardless of the ARID1A status (Fig. 2A). Importantly, BMAL2 depletion (Fig. 2B) inhibited cell proliferation (Fig. 2C), reduced DNA synthesis (Fig. 2D and S1A), and decreased mitosis (Fig. 2E and S1B) in both ARID1A-wt and ARID1A-mut cell lines. BMAL2 knockdown also inhibited clonogenic ability *in vitro* (Fig. 2F, S1C). Subcutaneous xenografts using NUDE mice injected with shCtrl or shBMAL2 (clone #1) lentiviral transduced ARID1A-wt OCCC cell lines, ES-2 and JHOC5, and ARID1A-mut OCCC cell line, OVISE, showed that BMAL2 depletion reduced tumor growth (Fig. 2G-2I). Immunohistochemistry (IHC) analysis further confirmed that shBMAL2 tumors had decreased proliferating cells indicated by reduced Ki67-positive cells (Fig. 2J-2L and S1D). Together, these results suggested that BMAL2 plays an important role in OCCC tumorigenesis.

**Figure 2.**
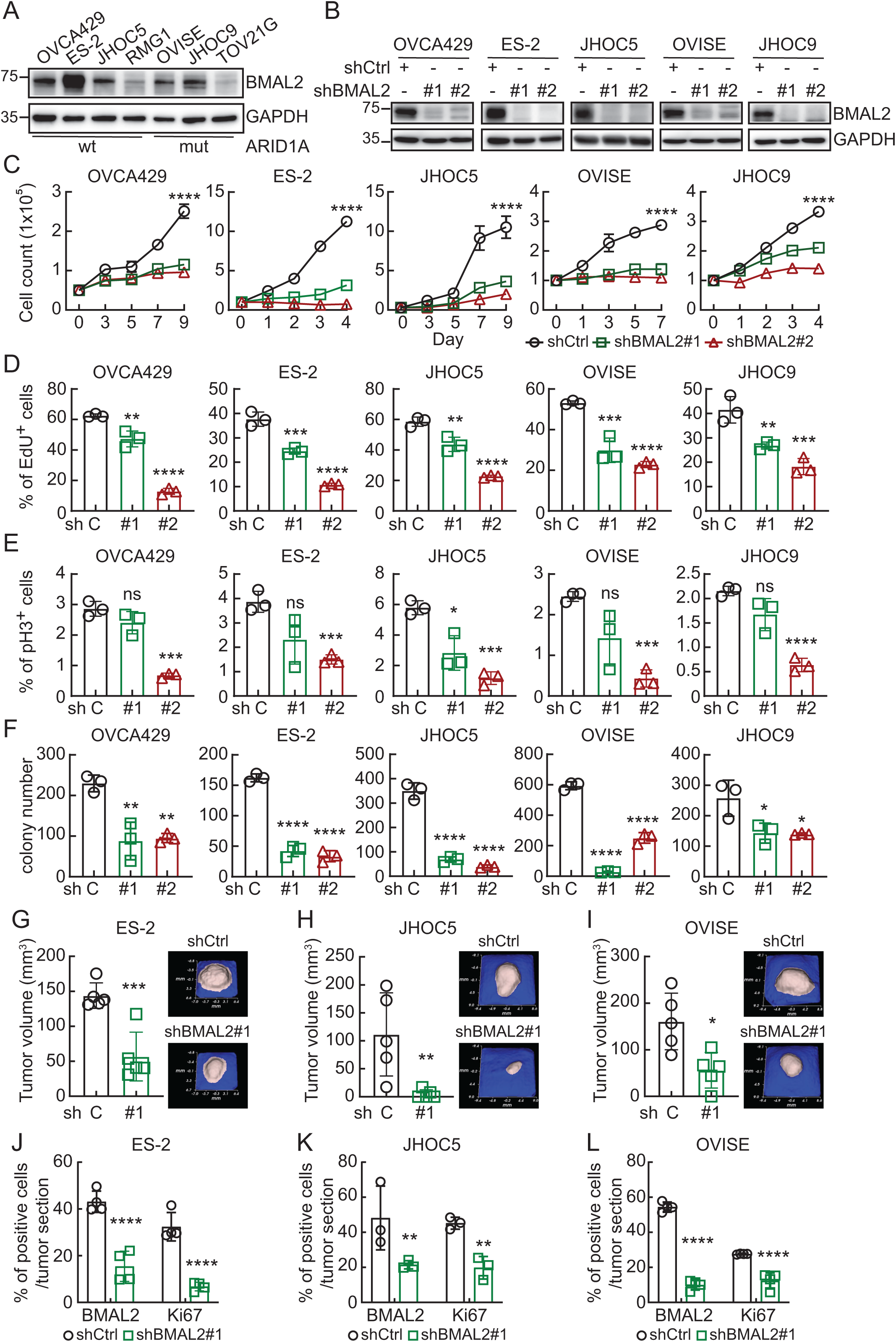
BMAL2 depletion inhibits tumorigenic ability in OCCC cells. **A.** IB of BMAL2 protein expression with GAPDH as a loading control. Blots shown are from one representative experiment of three replicates. **B.** IB of BMAL2 protein expression with GAPDH as a loading control in BMAL2-depleted (shBMAL2#1 or #2) OCCC cells. Blots shown are from one representative experiment of three replicates. **C.** Cell growth assays using control (shCtrl) or BMAL2-depleted (shBMAL2) OCCC cells. Data are shown as mean ± SD with *P* value based on two-way ANOVA test (n = 3). ****, *P* < 0.0001. The experiments were repeated 3 times. **D.** EdU staining assays using shCtrl or shBMAL2 OCCC cells. Data are shown as mean ± SD with *P* value based on unpaired *t* test (n = 3). **, *P* < 0.01; ***, *P* < 0.001; ****, *P* < 0.0001. The experiments were repeated 3 times. **E.** Phospho-histone H3 (pHH3) staining assays using shCtrl or shBMAL2 OCCC cells. Data are shown as mean ± SD with *P* value based on unpaired *t* test (n = 3). *, *P* < 0.05; ***, *P* < 0.001; ****, *P* < 0.0001; ns, not significant. The experiments were repeated 3 times. **F.** Clonogenic assays using shCtrl or shBMAL2 OCCC cells. Data are shown as mean ± SD with *P* value based on unpaired *t* test (n = 3). *, *P* < 0.05; ***, *P* < 0.001; ****, *P* < 0.0001. The experiments were repeated 3 times. **G, H and I.** Xenograft models in NUDE mice using shCtrl or shBMAL2 ES-2, JHOC5 and OVISE cells. Five mice were used for each group. Data are means ± SD, significant difference is based on unpaired *t* test of the tumor volume 3, 4 and 8 weeks after subcutaneous injection, respectively. *, *P* < 0.05; **, *P* < 0.01; ***, *P* < 0.001. **J, K and L.** Quantification and comparison of percentage of BMAL2 and Ki67 expressing cells detected by IHC in shCtrl or shBMAL2 ES-2, JHOC5 and OVISE derived tumors. Significant differences are based on unpaired *t* test. **, *P* < 0.01; ****, *P* < 0.0001.

### Loss of BMAL2 increases endogenous DNA damages in OCCC cells

To understand how BMAL2 depletion inhibited OCCC tumorigenesis, RNA-seq analysis was performed using ARID1A-wt OVCA429 and ARID1A-mut OVISE cells without or with BMAL2 depletion. KEGG pathway enrichment analysis indicated that differential expressed genes (DEG) were enriched in pathways including DNA damage repair (DDR), homologous recombination (HR), cell cycle and cellular senescence in both cell lines (Fig. 3A). These pathways are functionally interconnected as DDR and HR are activated upon DNA damage. And if the damage is too severe or cannot be fixed, the cell cycle can be arrested, leading to cellular senescence.

**Figure 3.**
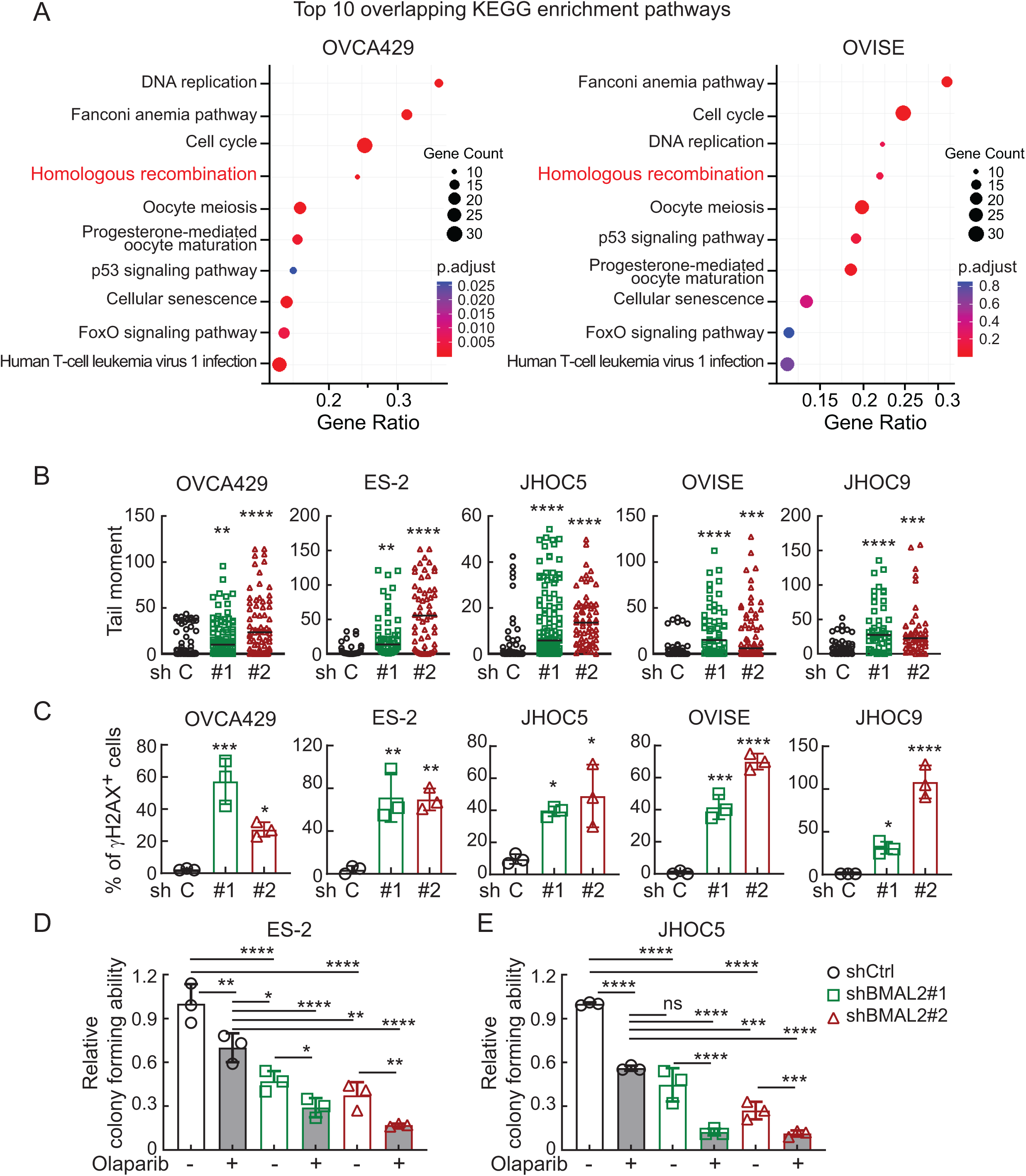
BMAL2 depletion increases endogenous DNA damages in OCCC cells. **A.** KEGG Enrichment Analysis of downregulated genes in BMAL2-depleted OVCA429 and OVISE cells. The top 10 overlapping pathways, determined by their adjusted *P* value, are shown. **B.** Tail moment of comet assays using shCtrl or shBMAL2 OCCC cells. At least 50 cells per group were measured. Significant differences are based on unpaired *t* test. **, *P* < 0.01; ***, *P* < 0.001; ****, *P* < 0.0001. The experiments were repeated 3 times. **C.** γH2AX staining assays using shCtrl or shBMAL2 OCCC cells. Data are shown as mean ± SD with *P* value based on unpaired *t* test (n = 3). *, *P* < 0.05; **, *P* < 0.01; ***, *P* < 0.001; ****, *P* < 0.0001. The experiments were repeated 3 times. **D.** Clonogenic assays using shCtrl or shBMAL2 OCCC cells without or with Olaparib treatment. Data are shown as mean ± SD with *P* value based on unpaired *t* test (n = 3). *, *P* < 0.05; **, *P* < 0.01; ***, *P* < 0.001; ****, *P* < 0.0001; ns, not significant. The experiments were repeated 3 times.

To validate whether BMAL2 depletion increased DNA damage, the physical state of DNA and biological response to DNA double-strand breaks were evaluated using alkaline comet assay (25) and γH2AX (26) staining, respectively. The comet assay showed a significant increase in tail moment in the BMAL2 depleted cells (Fig. 3B and S2A), suggesting that loss of BMAL2 leads to DNA damage. BMAL2 knockdown also elevated γH2AX level and foci numbers (Fig. 3C, S2B and S2C), indicating that BMAL2 depletion significantly increased DNA double-strand breaks. It is worth mentioning that the DNA damage resulted from BMAL2 depletion was only observed in OCCC cells, not in serous OC subtypes. BMAL2 knockdown did not lead to increase in γH2AX foci in HEYA8 and KURAMOCHI cells (Fig. S2D), suggesting that BMAL2-mediated DNA integrity maintenance is an OCCC subtype-specific regulatory mechanism.

Importantly, most of the genes in HR pathway were downregulated when BMAL2 was depleted (Fig. S3A, S3B and Table S4). In particular, expression of RAD51, a crucial protein responsible for the core steps of HR in eukaryotes (27), was significantly downregulated (Fig. S3C and S3D). Since BMAL2 is a transcriptional factor that regulates its downstream gene expression via binding to E-boxes, a regulatory motif of DNA with the consensus sequence 5’-CANNTG-3’ (28), we further investigate whether BMAL2 directly upregulated RAD51 transcription. We used The Eukaryotic Promoter Database (EPD) (29) to examine the RAD51 proximal promoter and found three E-box containing motifs (-983 to -978, -802 to 797, -209 to -202) (Fig. S3E). Chromatin immunoprecipitation (ChIP) further confirmed the binding of BMAL2 to these E-box regions (Fig. S3F), indicative of activation of RAD51 transcription when BMAL2 was present.

Since deficiency in RAD51 is known to impair HR and lead to DNA damage accumulation (27), and BMAL2 depletion resulted in downregulation of RAD51 (Fig. S3C and S3D), we employed an HR activity reporter assay (23) to determine whether BMAL2 depletion led to HR deficiency and therefore resulted in DNA damage accumulation. As a comparison, we treated the cells with B02, a RAD51 specific inhibitor (Fig. S3G). BMAL2 depletion greatly inhibited HR, even more so than B02 treatment (Fig. S3G). Collectively, RAD51 transcriptional regulation is one of the multifaceted functions of BMAL2 to maintain HR and prevent DNA damage in OCCC cells.

### Depletion of BMAL2 alone or with PARP inhibitor (PARPi) can inhibit tumorigenic ability of ARID1A-wt OCCC cells

Olaparib, a PARP inhibitor (PARPi), has been approved in the United States and Europe as a maintenance therapy for platinum-sensitive relapsed BRCA-mutated OC that displays defects in the HR repair pathway. It has been reported that 6.25 µM Olaparib can kill half of the population (IC_50_) of primary OC cells with HR deficiency (23). We examined the sensitivity of OCCC cells to Olaparib (dose range 0 to 10 µM) and found that ARID1A-mut OCCC cells were more sensitive to PARPi than ARID1A-wt OCCC cells (Fig. S4). The Olaprib IC_50_ was below 2.5 µM for OVISE (ARID1A-mut) cells, but was much higher for ARID1A-wt OCCC cells.

Since BMAL2 depletion impaired HR in OCCC, we reasoned that targeting BMAL2 may enhance PARPi sensitivity in ARID1A-wt OCCC cells and inhibit their tumorigenic ability. In the shCtrl OCCC cells, 2.5 µM Olaparib reduced colony numbers by 30% in ES-2 cells (Fig. 3D) and 40-45% in JHOC5 cells (Fig. 3E). Importantly, depletion of BMAL2 was itself able to achieve a much more pronounced inhibitory effect (50% reduction) than Olaparib in ES-2 cells (Fig. 3D). When BMAL2-depletion was combined with Olaparib, the colony forming ability of ARID1A-wt OCCC cells was further reduced (Fig. 3D and 3E). These results indicated that targeting BMAL2, alone or with PARPi, can be an effective therapeutic strategy for ARID1A-wt OCCC.

### GW833972A can be used to target BMAL2 for ARID1A-wt OCCC treatment

Transcription factors are traditionally thought to be undruggable; however, recent development of small-molecule modulators has suggested that targeting transcription factors can be feasible (30). We utilized a structure-based virtual screening to evaluate potential drugs that can be used to target BMAL2. Currently, the full-length BMAL2 protein structure has not been solved. The C-terminal PAS2 domain (N360-E477) of BMAL2 is the only available 3D structure resolved by NMR technique [Protein Database (PDB) Entry- 2KDK, Fig. S5A]. Thus, we used the PAS2 domain structure and Alphafold protein structure database-predicted BMAL2 full-length 3D structure to screen for potential binding pockets. A surface binding pocket containing E406, Y407, F408, H409, Q410, R438, A439 and F444 residues in the PAS2 domain was found and used as a docking area to perform virtual screening (Fig. S5B). Based on the docking scores, drug properties, and binding mode analysis, 119 high-affinity compounds (FRED Chemgauss4 score between -9.3 and -7.8) were identified. The top 9 bioactive compounds with low Kd value were then selected for further experiments. Impact of each selected compound on BMAL2 protein level was evaluated using ARID1A-wt ES-2 cells (Fig. S5C). GW833972A, a cannabinoid receptor 2 (CB2) selective agonist (31), was the most effective compound to reduce BMAL2 expression at the lowest dose (5 µM, Fig. S5C). Importantly, GW833972A did not downregulate BMAL2 mRNA expression in any of five OCCC cell lines tested (Fig. 4A), but substantially reduced its protein level in all five cell lines (Fig. 4B). To determine whether GW833972A binds to BMAL2 protein, cellular thermal shift assay (CETSA) and drug affinity responsive target stability (DARTS) analysis were performed. The thermal stability of BMAL2 protein was significantly increased (Fig. 4C) and BMAL2 protein was more resistant to enzymatic digestion when GW833972A was present (Fig. 4D). Cycloheximide (CHX) chase assay further demonstrated that BMAL2 protein was less stable when the cells were treated with GW833972A (Fig. 4E and 4F). Together, results from these biochemical assays suggested that GW833972A physically interacted with BMAL2 protein (32,33), as predicted in the structure-based virtual modeling (fig. S5D), and facilitated BMAL2 protein degradation.

**Figure 4.**
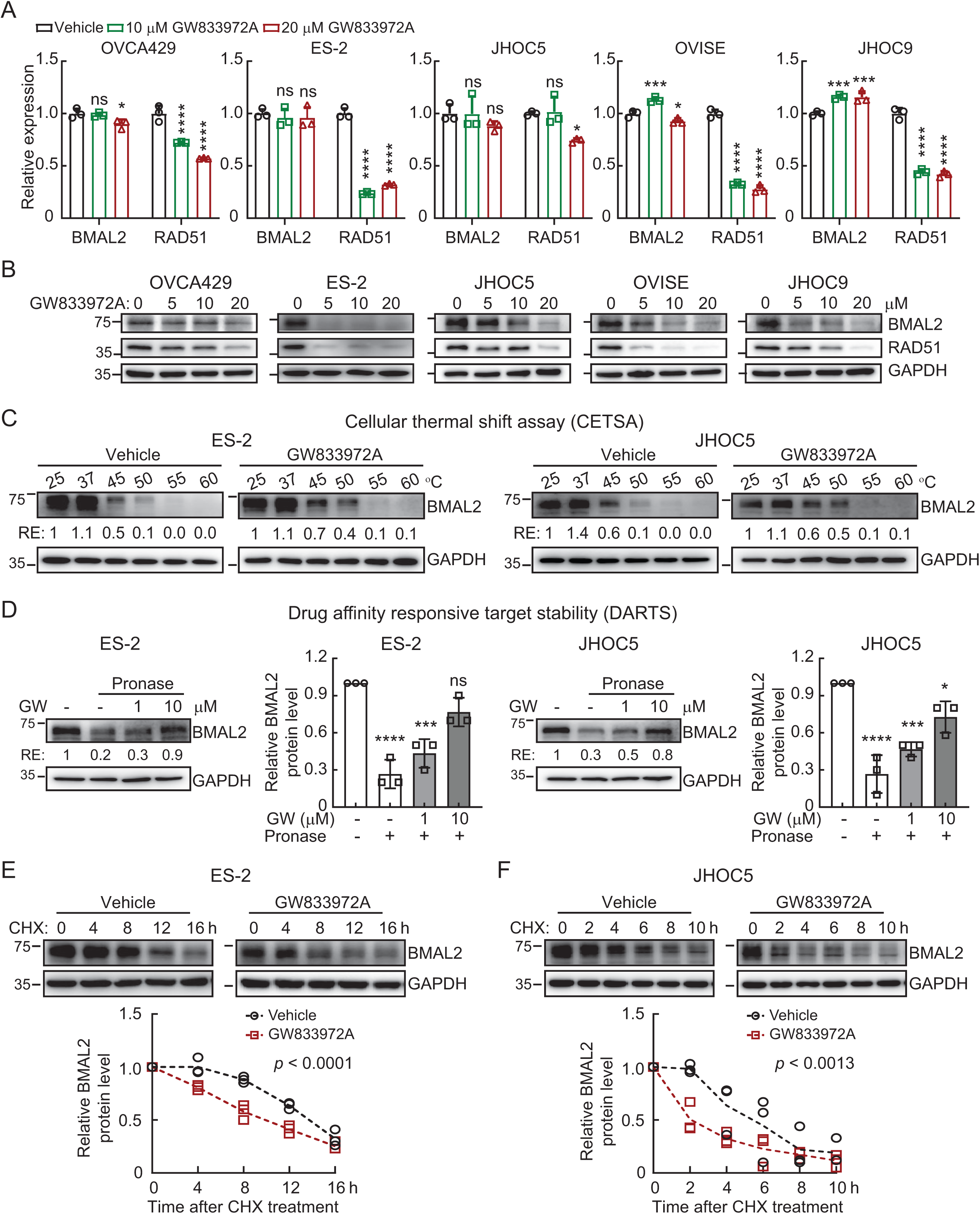
GW833972A targets BMAL2 protein for degradation. **A.** qRT-PCR analysis of BMAL2 and RAD51 level in OCCC cells treated with vehicle, 10 or 20 µM GW833972A. RNA18S5 was used as an internal control. Three independent experiments were performed and data are means ± SD from one representative experiment (n = 3). *, *P* < 0.05; ***, *P* < 0.001; ****, *P* < 0.0001; ns, not significant. Significant differences are based on unpaired *t* test. **B.** IB of BMAL2 and RAD51 protein expression with GAPDH as a loading control in OCCC cells treated with vehicle, 5, 10 or 20 µM GW833972A. Blots shown are from one representative experiment of three replicates. **C.** Cellular thermal shift assay (CETSA) using ES-2 and JHOC5 cells treated with vehicle or 20 µM GW833972A. The level of BMAL2 was determined using IB analysis. GAPDH was used as a loading control. Blots shown are from one representative experiment of three replicates. RE, relative expression. **D.** Drug affinity responsive target stability (DARTS) assay using ES-2 and JHOC5 cells treated with vehicle, 1 or 10 µM GW833972A, without or with 0.2 µg/ml Pronase. The level of BMAL2 was determined using IB analysis. GAPDH was used as a loading control. Blots shown are from one representative experiment of three replicates and relative BMAL2 protein levels from three IB analyses were quantified. Data are means ± SD. **E.** Time course assay using cycloheximide (CHX) treated ES-2 cells without or with 20 µM GW833972A treatment. The level of BMAL2 was determined using IB analysis. GAPDH was used as a loading control. Relative BMAL2 protein levels from three IB analyses were quantified and nonlinear regression (curve fit) analysis was used to test for significant differences between the degradation curve of the vehicle control and GW833972A treated group (*P* value is indicated). **F.** Time course assay using CHX treated JHOC5 cells without or with 20 µM GW833972A treatment. The level of BMAL2 was determined using IB analysis. GAPDH was used as a loading control. Data formatting is as described for E.

All five cell lines also showed reduced RAD51 expression after GW833972A treatment (Fig. 4A), indicative of decreased BMAL2 levels. GW833972A treatment also increased γH2AX foci (Fig. 5A), inhibited cell proliferation (Fig. 5B), decreased colony forming ability (Fig. 5C) and reduced cell viability (Fig. 5D) in a dose-dependent manner. All these phenotypes were consistent with the effect of decreasing BMAL2 protein levels by shRNA (Fig. 2 and 3). Also, GW833972A treatment significantly enhanced Olaparib sensitivity in ES-2 and JHOC5 cells with cell viability decreased from 75% to 40% (Fig. 5E and 5F). Importantly, a relatively high dose of GW833972A (20µM) alone reduced cell viability to 40% (Fig. 5E and 5F). At this high dose of GW833972A, combination with Olaparib did not further suppress cell viability (Fig. 5E and 5F). These results indicate that GW833972A is effective by itself at high dose and can also be used at lower dosages to enhance the effectiveness of PARPi treatments in ARID1A-wt OCCC.

**Figure 5.**
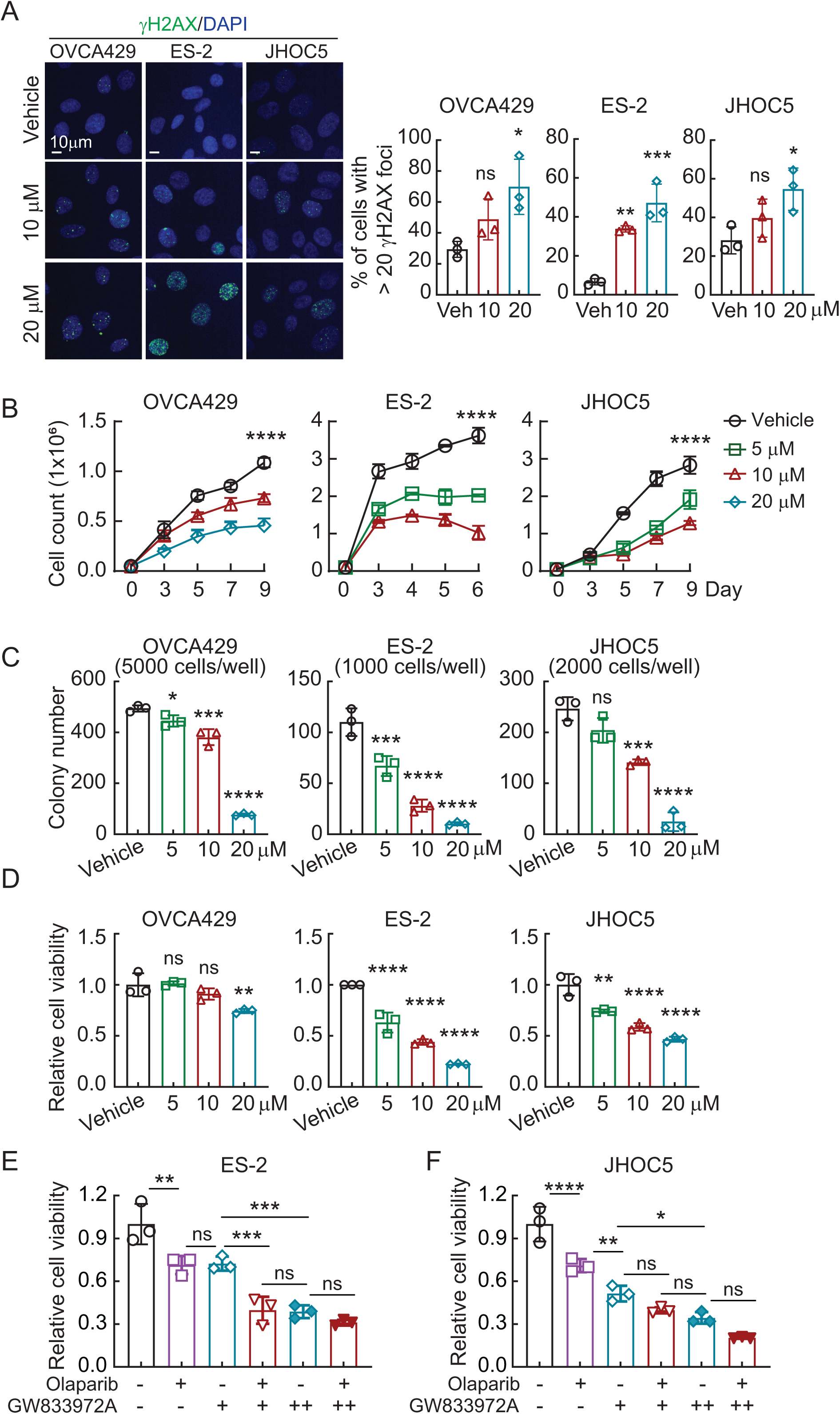
GW833972A inhibits tumorigenic ability in ARID1A-wt OCCC cells. **A.** Representative γH2AX immunofluorescent staining of ARID1A-wt OCCC cells treated with vehicle, 10 or 20 µM GW833972A. Percentage of cells with more than 20 γH2AX foci per nuclear was quantified. Data are shown as mean ± SD with *P* value based on unpaired *t* test (n = 3). *, *P* < 0.05; **, *P* < 0.01; ***, *P* < 0.001; ns, not significant. The experiments were repeated 3 times. **B.** Cell growth assays using ARID1A-wt OCCC cells treated with vehicle, 5, 10 or 20 µM GW833972A. Data are shown as mean ± SD with *P* value based on two-way ANOVA test (n = 3). ****, *P* < 0.0001. The experiments were repeated 3 times. **C.** Clonogenic assays using ARID1A-wt OCCC cells treated with vehicle, 5, 10 or 20 µM GW833972A. Data are shown as mean ± SD with *P* value based on unpaired *t* test (n = 3). *, *P* < 0.05; ***, *P* < 0.001; ****, *P* < 0.0001; ns, not significant. The experiments were repeated 3 times. **D.** Cell viability assays using ARID1A-wt OCCC cells treated with vehicle, 5, 10 or 20 µM GW833972A. Data are shown as mean ± SD with *P* value based on unpaired *t* test (n = 3). **, P < 0.01; ****, P < 0.0001; ns, not significant. The experiments were repeated 3 times. **E.** Cell viability assays using ES-2 cells treated with vehicle, 10 (+) or 20 (++) µM GW833972A with vehicle, 5 (+) or 10 (++) µM Olaparib. Data formatting is as described for D. **, *P* < 0.01; ***, *P* < 0.001; ****, *P* < 0.0001; ns, not significant. **F.** Cell viability assays using JHOC5 cells treated with vehicle, 10 (+) or 20 (++) µM GW833972A with vehicle, 2.5 (+) or 5 (++) µM Olaparib. Data formatting is as described for D. *, *P* < 0.05; **, *P* < 0.01; ****, *P* < 0.0001; ns, not significant.

To determine whether GW833972A is toxic to normal cells, WI38 human fetal lung fibroblasts were used. A slight decrease in cell viability (Fig. S6A), but no DNA damage (Fig. S6B), was observed when WI38 cells were treated with 20 μM GW833972A. Moreover, GW833972A had a much higher IC_50_ for WI38 fibroblasts (IC_50_: 133.6 μM) than for ES-2 OCCC cells (IC_50_: 20.2 μM) (Fig. S6C). The low toxicity of GW833972A in WI38 fibroblasts was likely due to the extremely low BMAL2 expression, further demonstrating the drug-target specificity between GW833972A and BMAL2 (Fig. 4C-4F and S6D). Since BMAL2 expression is low in most normal tissues (Fig. 1A), these observations suggest that GW833972A can effectively target BMAL2 expressing cancer cells with little adverse effects on normal cells.

To validate these cell-based observations in vivo, subcutaneous xenograft models using ES-2 and JHOC5 cells were performed (Fig. 6 and S7). When tumors reached 50 mm^3^, mice were intraperitoneally injected with either vehicle or 10 mg/kg GW833972A daily for five days a week over the course of two weeks (Fig. S7A). The experiments were terminated when tumor-induced skin breakage occurred in the vehicle control group. In both mouse models, GW833972A treatment significantly inhibited tumor growth without body weight changes (Fig. 6A, 6B, S7B and S7C). No abnormalities were observed in the liver, kidney and spleen in these mice (Fig. S7D and S7E), suggesting low adverse side-effects of GW833972A. These tumors showed reduced BMAL2 and RAD51 positive cells, increased DNA double- stranded breaks (γH2AX) and decreased cell proliferation (Ki67) compared to the vehicle treated control (Fig. 6C-6F). Together, the data showed that targeting BMAL2 by GW833972A provides a therapeutic opportunity for ARID1A-wt OCCC.

**Figure 6.**
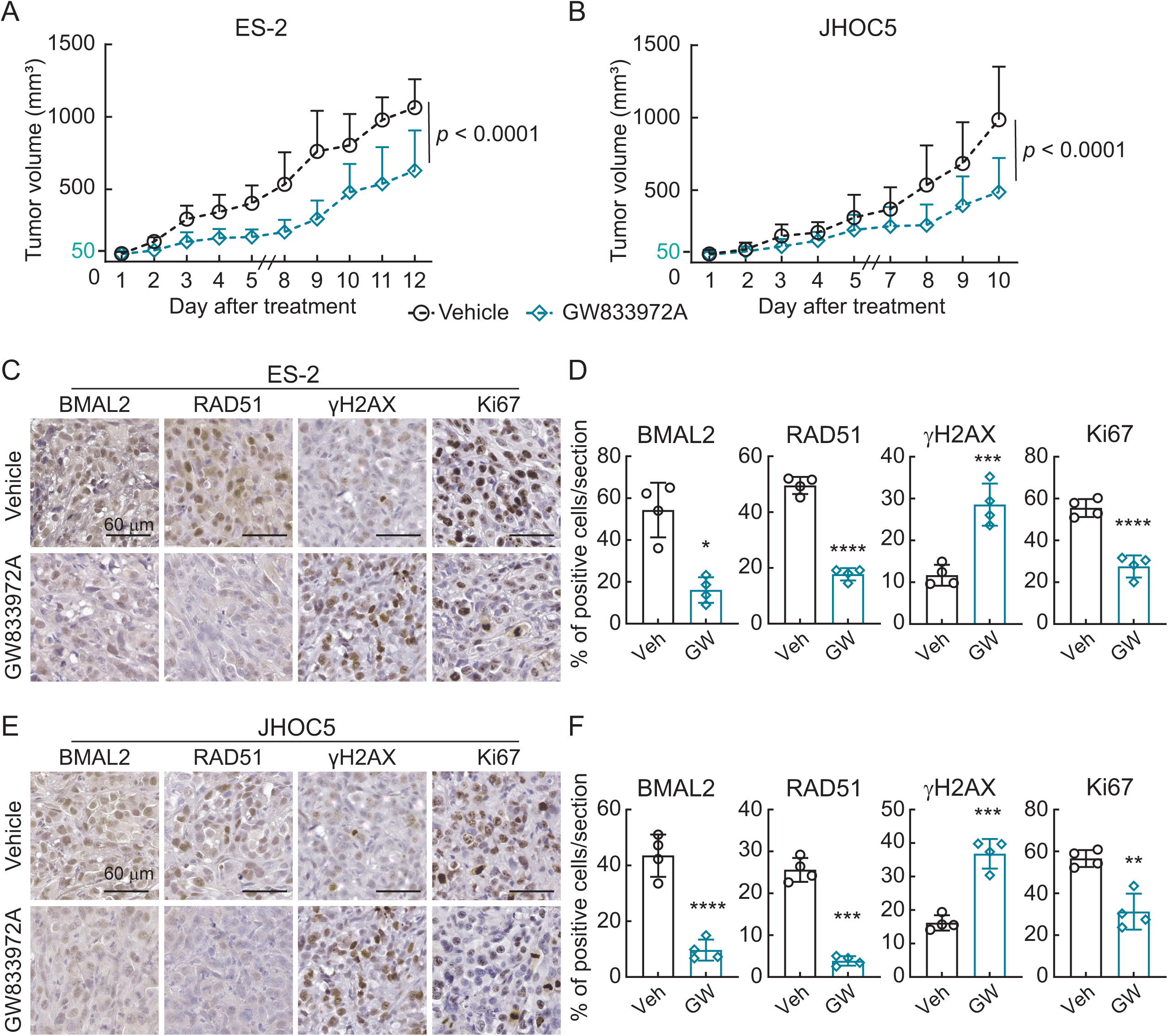
Targeting BMAL2 by GW833972A increases DNA damages and inhibits ARID1A-wt OCCC tumor growth. **A.** Subcutaneous xenograft model in NUDE mice using ES-2 cells with or without 10 mg/kg GW833972A treatment. Five mice were used for each group. Nonlinear regression (curve fit) analysis was used to test for significant differences between the tumor growth curve of the vehicle control and GW833972A treated group (*P* value is indicated). **B.** Subcutaneous xenograft model in NUDE mice using JHOC5 cells with or without 10 mg/kg GW833972A treatment. Five mice were used for each group. Data formatting is as described for A. **C and D.** Representative images of BMAL2, RAD51, γH2AX and Ki67 IHC staining using serial tumor sections from ES-2 derived tumors (C). (Scale bars, 60 μm.) Percentage of positive cells was quantified (D). Data are shown as mean ± SD with *P* value based on unpaired *t* test. **, *P* < 0.01; ***, *P* < 0.001; ****, *P* < 0.0001; ns, not significant. **E and F.** Representative images of BMAL2, RAD51, γH2AX and Ki67 IHC staining using serial tumor sections from JHOC5 derived tumors (E). (Scale bars, 60 μm.) Percentage of positive cells was quantified (F). Data formatting is as described for D.

## Discussion

Current therapeutic strategies for OC are based on the results of clinical trials for HGSOC. Unfortunately, OCCC is relatively insensitive to the standard carboplatin-paclitaxel adjuvant chemotherapy, and due to its distinct molecular features often becomes more malignant with worse prognosis at the advanced stage compared to other OC subtypes (11). Therapeutic strategies, such as immune checkpoint blockade, have been tested but failed to deliver promising results for OCCC (14). It has been suggested that OCCC progression may partially depend on angiogenesis (11). Therefore, targeting the angiogenesis pathway was expected to deliver a better outcome. However, the enrolled OCCC patients showed no significant benefit from anti-angiogenesis drug bevacizumab (1). Also, use of other angiogenesis inhibitors as a single maintenance therapy for OCCC, such as Sunitinib and Nintedanib, which target platelet derived growth factor receptor (PDGFR) and multiple tyrosine kinases in the angiogenesis pathway, respectively, did not show high response rates (14). It is apparent that new targeted therapies and/or combination treatment strategies specifically for OCCC is urgently required.

ARID1A acts as a tumor suppressor and participates in DNA damage repair process (34). Thus, loss of ARID1A will increase sensitivity to DNA damaging agents (35). Many preclinical studies have utilized molecular mechanistic results and found ARID1A synthetic lethal partners, including PARP (36), Ataxia telangiectasia and Rad3-related (ATR) (37), Aurora kinase A (AURKA) (38), YES1 (39), Polycomb repressive complex 2 (PRC2) component EZH2 (40), and factors in glutathione metabolism pathway (41). In these studies, inhibition of the functions or suppression of the expression of these factors increased cell death in ARID1A deficient cancer cells. However, in clinical trials, the response rate of ARID1A-mut OCCC patients was disappointing (14,42). A few clinical trials using combination therapies are on-going (42) and the outcomes are uncertain.

Thus, it is evident that no promising targeted therapy has yet been developed for patients with ARID1A-wt OCCC. Based on our findings that BMAL2 expression was extremely low in non-tumor cells in the ovarian tissues but upregulated in OCCC cells, and that targeting BMAL2 led to excessive DNA damage, reduced cell viability and inhibited tumor growth, we proposed that BMAL2 can serve as a biomarker as well as a key therapeutic target, particularly for ARID1A-wt OCCC. Moreover, since BMAL2 regulates DNA damage repair pathways to prevent DNA damage, its upregulation in OCCC may also be an important contributing factor to the high chemoresistance of this refractory disease.

In mammalian cells, the core clock gene BMAL1 has also been shown to regulate DNA damage repair pathway (43). Although BMAL2 has structural and functional similarities to BMAL1 (44), its interaction and regulation with other central clock genes are different from that of BMAL1 in normal tissues (45). The exact role of BMAL2 in the circadian system is not yet been fully understood. It has been shown that transgenic overexpression of BMAL2 can rescue the loss of behavioral rhythms in *Bmal1* knockout mice (46). Nevertheless, such compensatory function is not observed naturally because *Bmal2* expression is decreased in *Bmal1* knockout mice (28,45). In contrast, BMAL2 indirectly suppresses BMAL1 expression via REV-ERBα/β and therefore depletion of BMAL2 can lead to upregulation of BMAL1 in non-cancerous cell lines (28). In the context of OC, BMAL2 is upregulated (Fig. 1) while most of the core clock genes, including BMAL1, are downregulated compared to normal tissues (47). Importantly, when BMAL2 was depleted in OCCC cells, the level of BMAL1 did not increase (Fig. S8A), indicating that the regulatory mechanisms of the circadian system are disrupted in cancer cells and that BMAL2 oncogenic function may be circadian independent.

We further demonstrated that the small molecule GW833972A can induced BMAL2 protein degradation. GW833972A monotherapy or combination with PARPi significantly reduced cancer cell viability and impeded tumor growth. These findings revealed a new opportunity for ARID1A-wt OCCC patients. Lead modification and optimization of GW833972A and detailed pharmacokinetic analyses are worthy of further investigation. Furthermore, such targeting of BMAL2 can be applied more broadly to other BMAL2 upregulated cancers. For example, BMAL2 expression is also associated with poor prognosis in lung cancer, pancreatic cancer, acute myeloid leukemia (AML) and myeloma (Fig. S8B-S8E), indicating a pervasive role of BMAL2 in cancer malignancy. Although BMAL2 may carry out its oncogenic functions differently in different tissues, our observations that GW833972A elicits BMAL2 degradation, rather than affecting BMAL2 interaction with one or more other proteins, suggest that GW833972A may be broadly applicable to BMAL2-upregulated cancers even if the mechanism of BMAL2 action differs among those cancers. Both the underlying molecular mechanisms of BMAL2- mediated tumorigenesis and the possibility of using BMAL2 as a multi-cancer therapeutic target are promising areas for further work.

## Acknowledgement

This work was supported by Academia Sinica (AS) [AS-GCS-111-L01] and Taiwan National Science and Technology Council [NSTC-114-2320-B-001-016-MY3] to Wendy W. Hwang-Verslues. The authors would like to thank the following core facilities at Academia Sinica: the National RNAi Core Facility for providing shRNA reagents and related services; the Bioinformatics Core at Institute of Molecular Biology for providing the RNA-seq analysis services; the DNA Sequencing Core Facility of the Institute of Biomedical Sciences [funded by Academia Sinica Core Facility and Innovative Instrument Project (AS-CFII-113-A12)] for DNA sequencing analysis; the Advanced Optics Microscope Core Facility (funded by AS-CFII-114-A3) for microscope imaging technical support, and Academia Sinica SPF Animal Facility (AS-CFII-113-A7) for providing animal support.

