## Supplemental Tables and Figures for "BMAL2 is a druggable target for ARID1A-wildtype ovarian clear cell carcinoma (OCCC)"

| Table S1. Short tandem repeat of OCCC cell lines and human fetal lung fibroblast cells | | | | | | | | |
|---|---|---|---|---|---|---|---|---|
| **STR/Cell line** | **OVCA429** | **ES-2** | **JHOC5** | **OVISE** | **JHOC9** | **TOV21G** | **RMG1** | **WI38** |
| **TH01** | 9,9 | 9.3, 9.3 | 7,7 | 9,9.3 | 6,9 | 7,9.3 | 11,11 | 8,9.3 |
| **D5S818** | 11,12 | 11,13 | 10,10 | 10,10 | 10,10 | 12,13 | 12,12 | 10,10 |
| **D13S317** | 7,12 | 11,11 | 9,12 | 11,12 | 10,10 | 11,12 | 12,12 | 11,11 |
| **D7820** | 11,12 | 11,11 | 12,12 | 11,12 | 8,12 | 12,12 | 11,11 | 9,11 |
| **D165539** | 12,12 | 11,13 | 11,13 | 9,9 | 12,12 | 10,12 | 9,10 | 11,12 |
| **CSF1PO** | 12,13 | 10,15 | 10,12 | 9,11 | 10,13 | 13,15 | 10,10 | 10,12 |
| **Amelogenin** | X,X | X,X | X,X | X,X | X,X | X,X | X,X | X,X |
| **vWA** | 16,18 | 16,17 | 14,16 | 18,18 | 14,17 | 17,17 | 17,18 | 19,20 |
| **TPOX** | 9,11 | 8,12 | 11,11 | 8,8 | 8,11 | 8,11 | 11,11 | 8,8 |
| **Match to Test Sample** | 100% | 100% | 100% | 100% | 96% | 100% | 95.65% | 100% |
| **Database** | ExPASy | ExPASy | ExPASy | ExPASy | ExPASy | ExPASy | ExPASy | ExPASy |

| Table S2. Primers for qRT-PCR | | |
|---|---|---|
| **Gene** | **Forward (5'- 3')** | **Reverse (5'- 3')** |
| RNA18S5 | gtaacccgttgaaccccatt | ccatccaatcggtagtagcg |
| BMAL2 | agctgttggtcttgtccctg | caagtggctcctgcgatg |
| RAD51 | tctctggcagtgatgtcctgga | taaagggcggtggcactgtcta |

| Table S3. Primers for ChIP-qPCR | | |
|---|---|---|
| **Promoter site** | **Forward (5'- 3')** | **Reverse (5'- 3')** |
| E-box 1 | cgattctcatgcctcagcct | ataaacctggccaacgtggt |
| E-box 2 | ttactggcgtgaaccaccg | agaggaagggggcattgaat |
| E-box 3 | gatactctcgcctcggcctc | tacagactgccctcttccct |

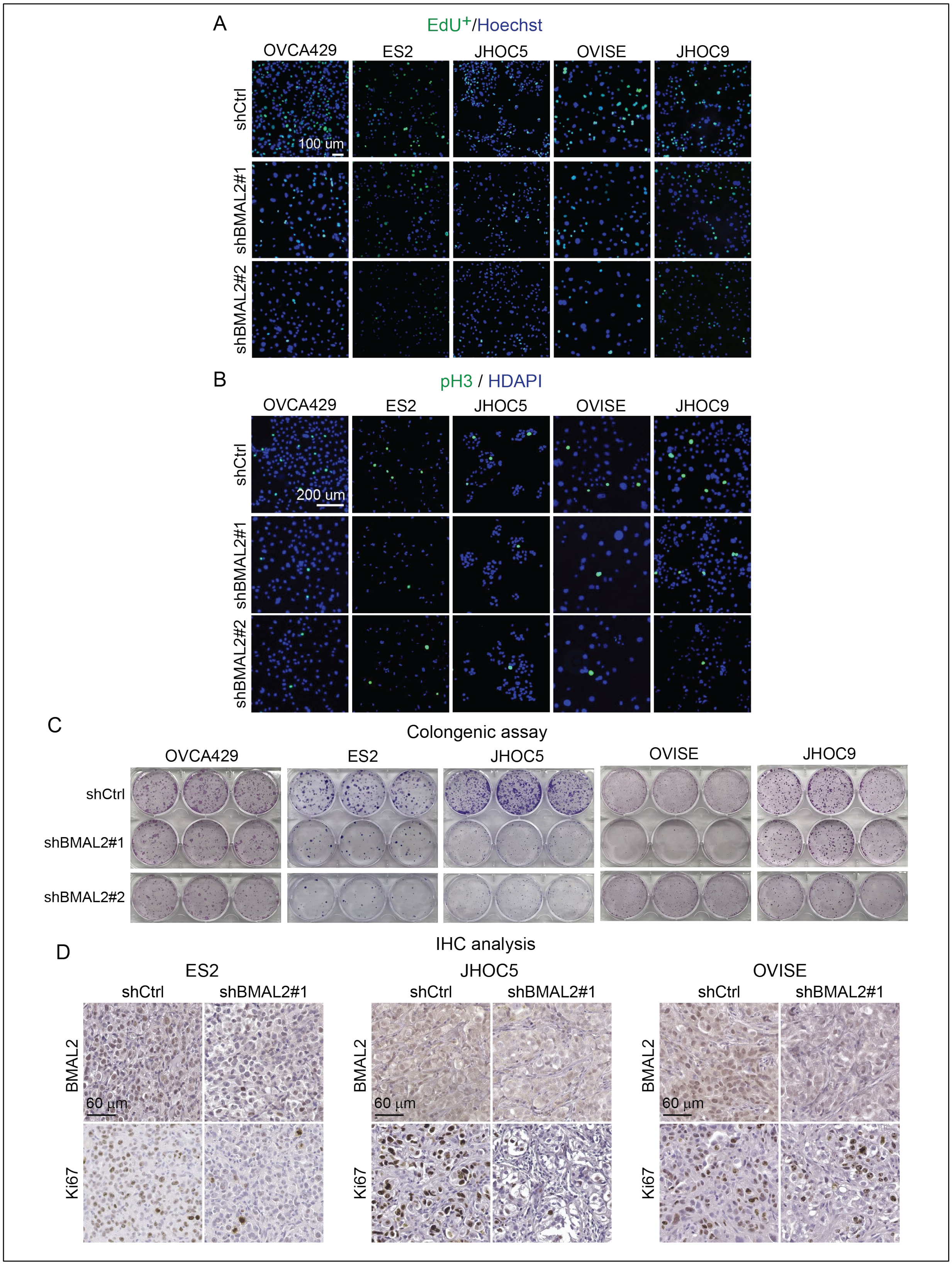

**Figure S1. BMAL2 depletion inhibits tumorigenic ability in OCCC cells.**

(A) Representative EdU staining of OCCC cells without (shCtrl) or with BMAL2 depletion (shBMAL2#1 or #2). Scale bar indicates 100 μm.

(B) Representative phospho-histone H3 (pHH3) staining of shCtrl and shBMAL2 OCCC cells. Scale bar indicates 200 μm.

(C) Representative images of clonogenic assays in OCCC cells.

(D) Representative images of BMAL2 and Ki67 IHC staining using serial tumor sections from ES-2, JHOC5 or OVISE derived tumors. Scale bars indicate 60 μm.

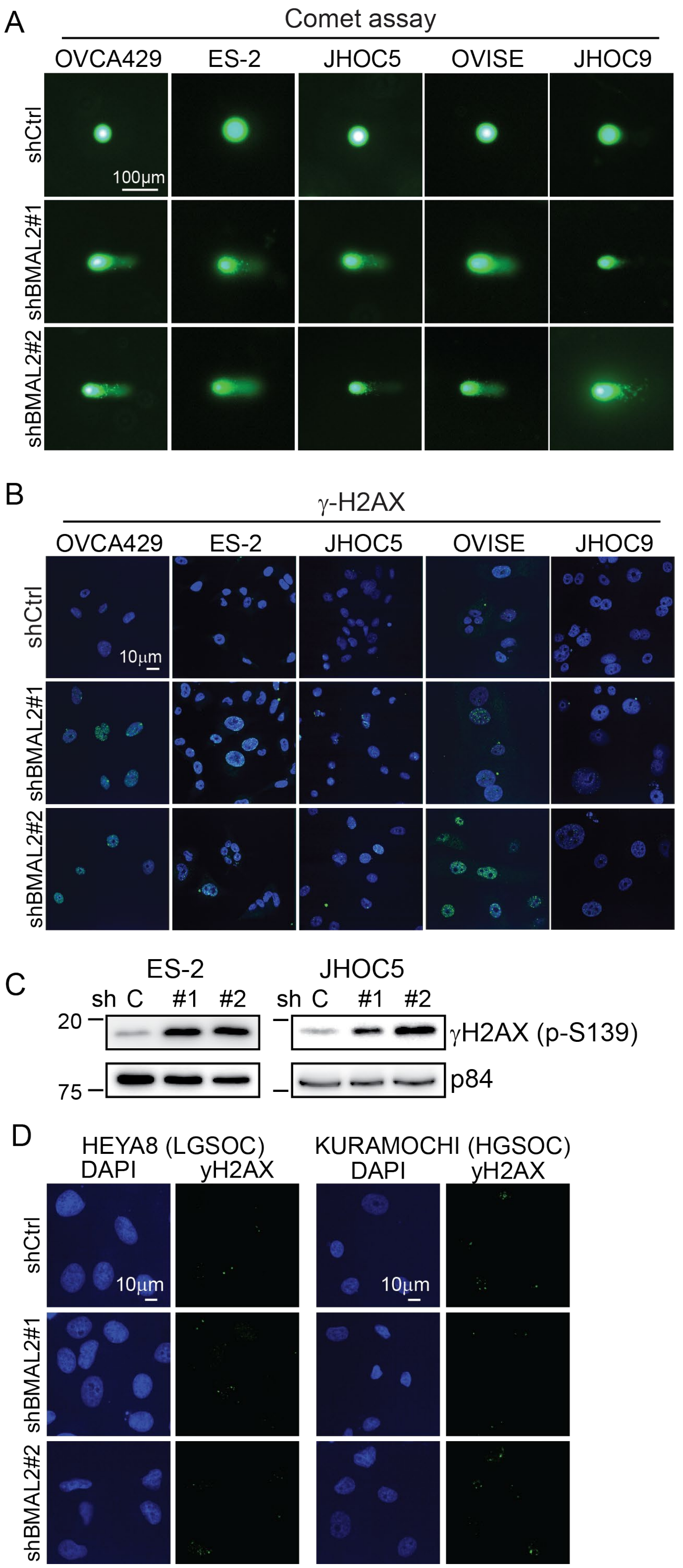

**Figure S2. BMAL2 depletion increases endogenous DNA damages in OCCC cells, but not serous type ovarian cancers**.

(A)  Representative comet assay images of OCCC cells without (shCtrl) or with BMAL2 depletion (shBMAL2#1 or #2).  Scale bar indicates 100 μm.

(B) Representative γH2AX staining of shCtrl and shBMAL2 OCCC cells.  Scale bar indicates 10 μm.

(C.) IB of γH2AX protein with p84 as a nuclear protein loading control.  Blots shown are from one representative experiment of three replicates.

(D) Representative γH2AX staining of shCtrl and shBMAL2 serous type OC cells.  Scale bars indicate 10 μm.

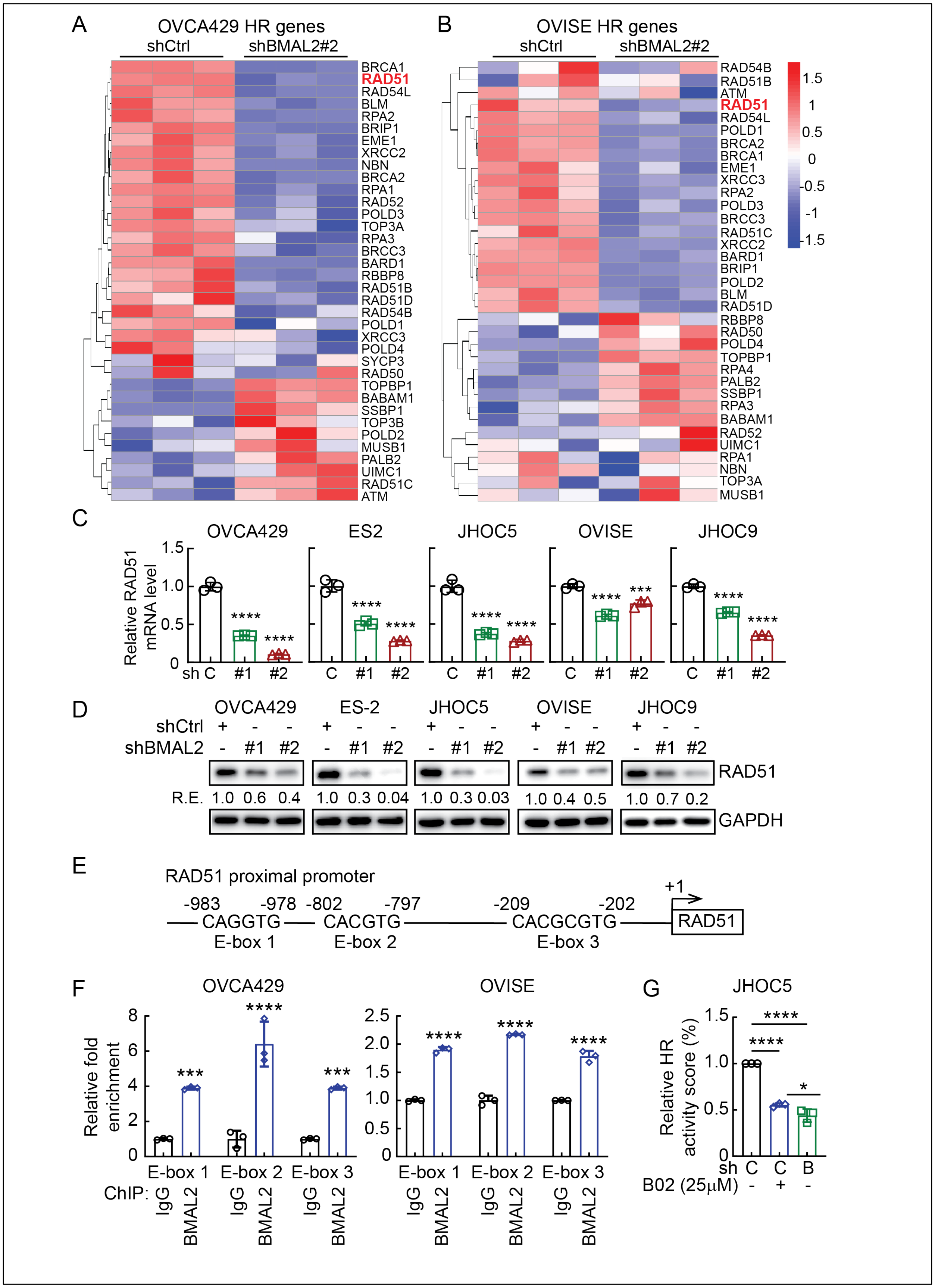

A
OVCA429 HR genes
shCtrl
shBMAL2#2
BRCA1
RAD51
RAD54L
BLM
RPA2
BRIP1
EME1
XRCC2
NBN
BRCA2
RPA1
RAD52
POLD3
TOP3A
RPA3
BRCC3
BARD1
RBBP8
RAD51B
RAD51D
RAD54B
POLD1
XRCC3
POLD4
SYCP3
RAD50
TOPBP1
BABAM1
SSBP1
TOP3B
POLD2
MUSB1
PALB2
UIMC1
RAD51C
ATM
B
OVISE HR genes
shCtrl
shBMAL2#2
RAD54B
RAD51B
ATM
RAD51
RAD54L
POLD1
BRCA2
BRCA1
EME1
XRCC3
RPA2
POLD3
BRCC3
RAD51C
XRCC2
BARD1
BRIP1
POLD2
BLM
RAD51D
RBBP8
RAD50
POLD4
TOPBP1
RPA4
PALB2
SSBP1
RPA3
BABAM1
RAD52
UIMC1
RPA1
NBN
TOP3A
MUSB1
1.5
1
0.5
0
-0.5
-1
-1.5
C
OVCA429
ES2
JHOC5
OVISE
JHOC9
Relative RAD51 mRNA level
1.5
1.0
0.5
0
****
***
sh C #1 #2
D
OVCA429
ES-2
JHOC5
OVISE
JHOC9
shCtrl + - -
shBMAL2 - #1 #2
RAD51
R.E. 1.0 0.6 0.4 1.0 0.3 0.04 1.0 0.3 0.03 1.0 0.4 0.5 1.0 0.7 0.2
GAPDH
E
RAD51 proximal promoter
+1
-983 -978 -802 -797 -209 -202
CAGGTG CACGTG CACGCGTG RAD51
E-box 1 E-box 2 E-box 3
F
OVCA429
OVISE
Relative fold enrichment
8
6
4
2
0
2.5
2.0
1.5
1.0
0.5
0
***
****
E-box 1 E-box 2 E-box 3
ChIP: IgG BMAL2
G
JHOC5
Relative HR activity score (%)
1.5
1.0
0.5
0
****
*
sh C C B
B02 (25μM) - + -

**Figure S3. BMAL2 depletion downregulates genes in homologous recombination (HR) pathway, including RAD51.**

(A) and (B) Heatmap of HR genes of the KEGG pathway (hsa03440) in OVCA429 (A) and OVISE (B) cells without or with BMAL2 depletion.

(C) qRT-PCR analysis of RAD51 level in OCCC cells without (shCtrl) or with BMAL2 depletion (shBMAL2#1 or #2). RNA18S5 was used as an internal control. Three independent experiments were performed and data are means ± SD from one representative experiment (n = 3). ***, $P < 0.001$; ****, $P < 0.0001$. Significant differences are based on unpaired *t* test.

(D) IB of RAD51 protein expression with GAPDH as a loading control in shCtrl or shBMAL2 OCCC cells. Blots shown are from one representative experiment of three replicates. RE, relative expression.

(E) Diagram shows three putative E-boxes on the *RAD51* promoter predicted using EPD eukaryotic promoter database.

(F) ChIP-qPCR analysis of BMAL2 on the *RAD51* promoter E-box regions. Three independent experiments were performed, and data are means ± SD from one representative experiment with significant differences detected by unpaired *t* test. ***, $P < 0.001$; ****, $P < 0.0001$.

(G) HR activity of shCtrl or shBMAL2 JHOC5 cells, assessed 72 h after adenovirus infection.

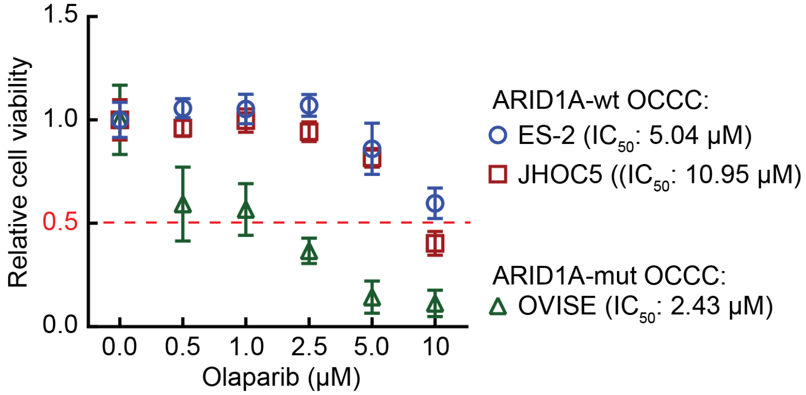

**Figure S4. ARID1A-mut OCCC cells are more sensitive to PARPi than ARID1A-wt OCCC cells.** Cell viability assays using ES-2, JHOC5 and OVISE cells treated with vehicle (0), 0.5, 1, 2.5, 5 or 10 µM Olaparib. Data are shown as mean ± SD (n = 3). The half-maximal inhibitory concentration ($IC_{50}$) of Olaparib for each cell line is indicated.

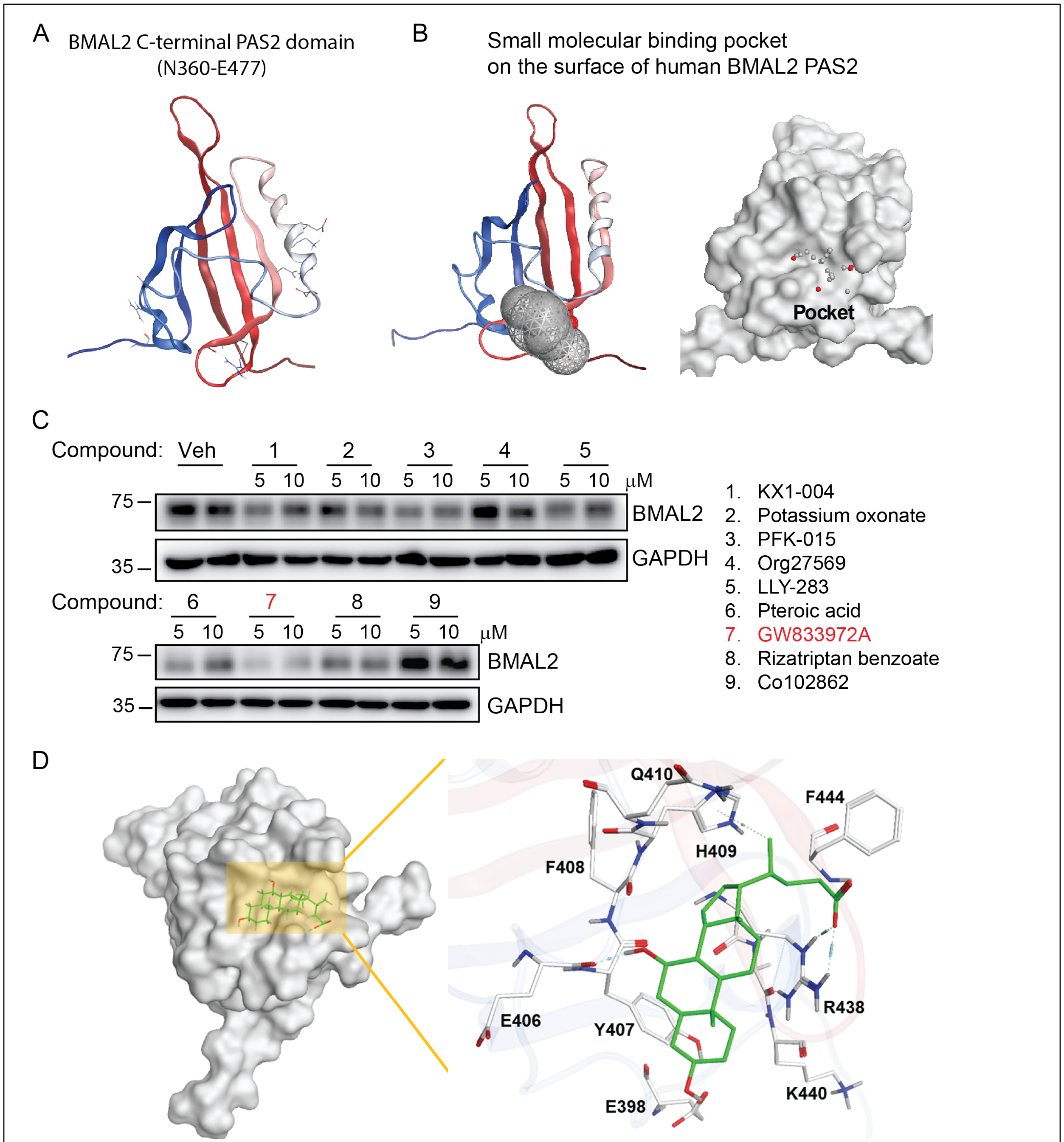

Left panel: BMAL2-GW833972A complex, in which the BMAL2 PAS2 is shown as molecular surface and GW833972A is shown in green.
Right panel: Interaction of T64394 to the binding area of BMAL2, where the contact residues were shown. Residue E406 formed hydrogen bonds and H409 formed π-hydrogen interaction with a ring of GW833972A.

**Figure S5. Virtual screening of potential small compounds for BMAL2 inhibition.**

(A) 3D structure of human BMAL2 PAS2 domain (2KDK). The N-terminal to C-terminal structure was colored from blue to red.

(B) The small molecular binding pocket on the surface of human BMAL2 PAS2 domain was identified using MOE-SiteFinder where the pocket was served as the docking areas for the virtual screening. The docking area was defined by a docking box with the length, width and height of 19.67 Å x 20.33 Å x 16.67 Å respectively and the total volume was 6664 $Å^3$, where the inner contour volume was 2614 $Å^3$.

(C) Left panel: IB of BMAL2 protein with GAPDH as a loading control in ES-2 cells treated with vehicle (DMSO), 5 or 10 μM selected compounds. Blots shown are from one representative experiment of three replicates. Right panel: The top 9 bioactive compounds with low Kd value used for evaluation.

(D) Binding modes of GW833972A to human BMAL2. BMAL2-compound complex, in which the BMAL2 PAS2 domain was shown as molecular surface and GW833972A was shown in green. Interaction of GW833972A to the contact residues of BMAL2 were shown.

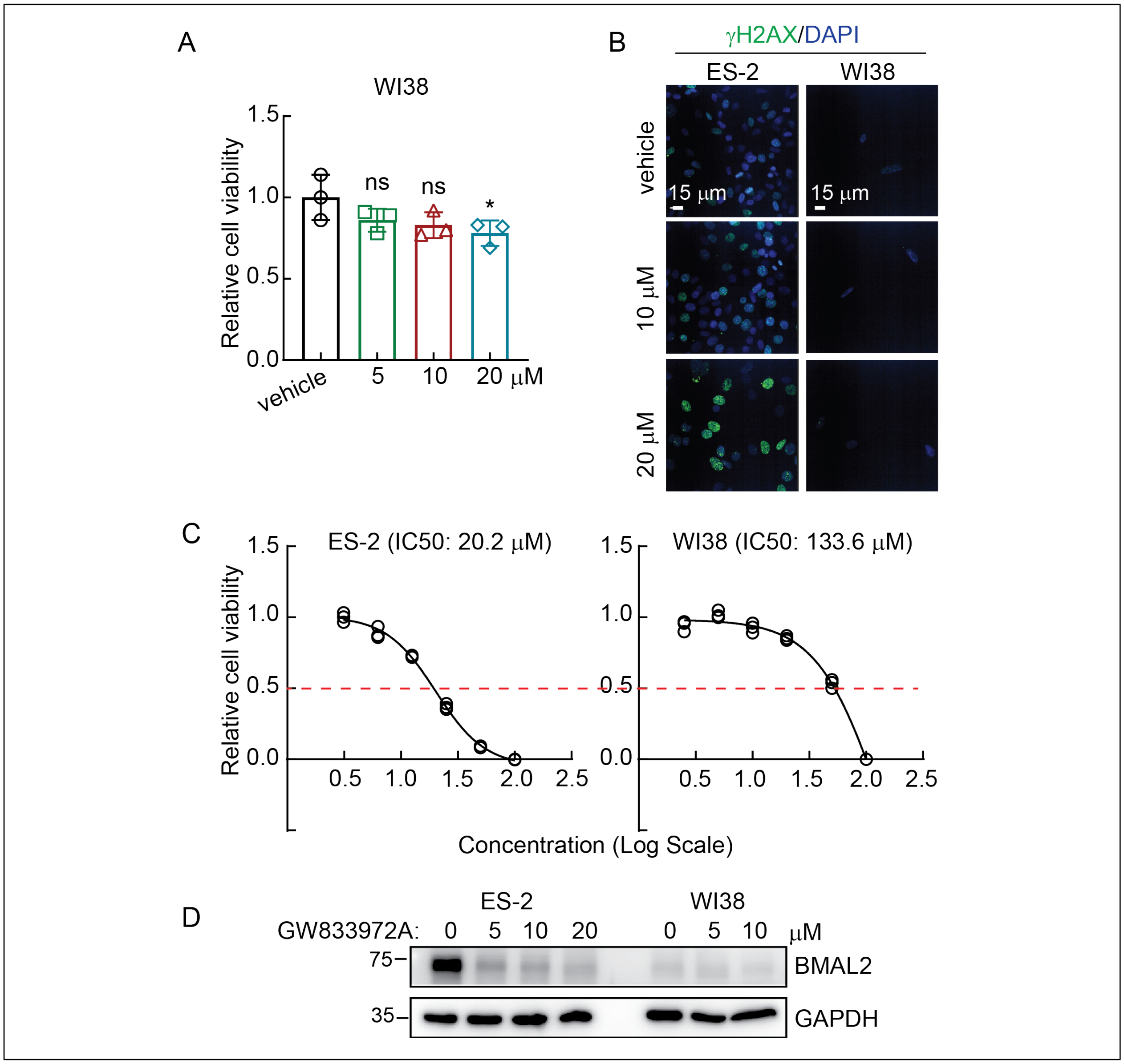

**Figure S6. GW833972A can effectively target BMAL2 expressing cancer cells while has low adverse effects on normal cells.**

(A) Cell viability assays using WI38 cells treated with vehicle (DMSO), 5, 10 or 20 µM GW833972A. Data are shown as mean ± SD with *P* value based on unpaired *t* test (n = 3). *, $P < 0.05$; ns, not significant. The experiments were repeated 3 times.

(B) Representative γH2AX staining of ES-2 and WI38 cells treated with vehicle (DMSO), 10 or 20 µM GW833972A. Scale bar indicates 15 µm.

(C) Dose-response curves for the assessment of cell viability in ES-2 and WI38 cells treated by GW833972A from 0 to 100 μM. The curves were plotted with log10 [GW833972A (μM)]. IC50 for each cell line was indicated.

(D) IB of BMAL2 protein with GAPDH as a loading control in ES-2 and WI38 cells treated with vehicle (DMSO), 5, 10 or 20 μM GW833972A. Blots shown are from one representative experiment of three replicates.

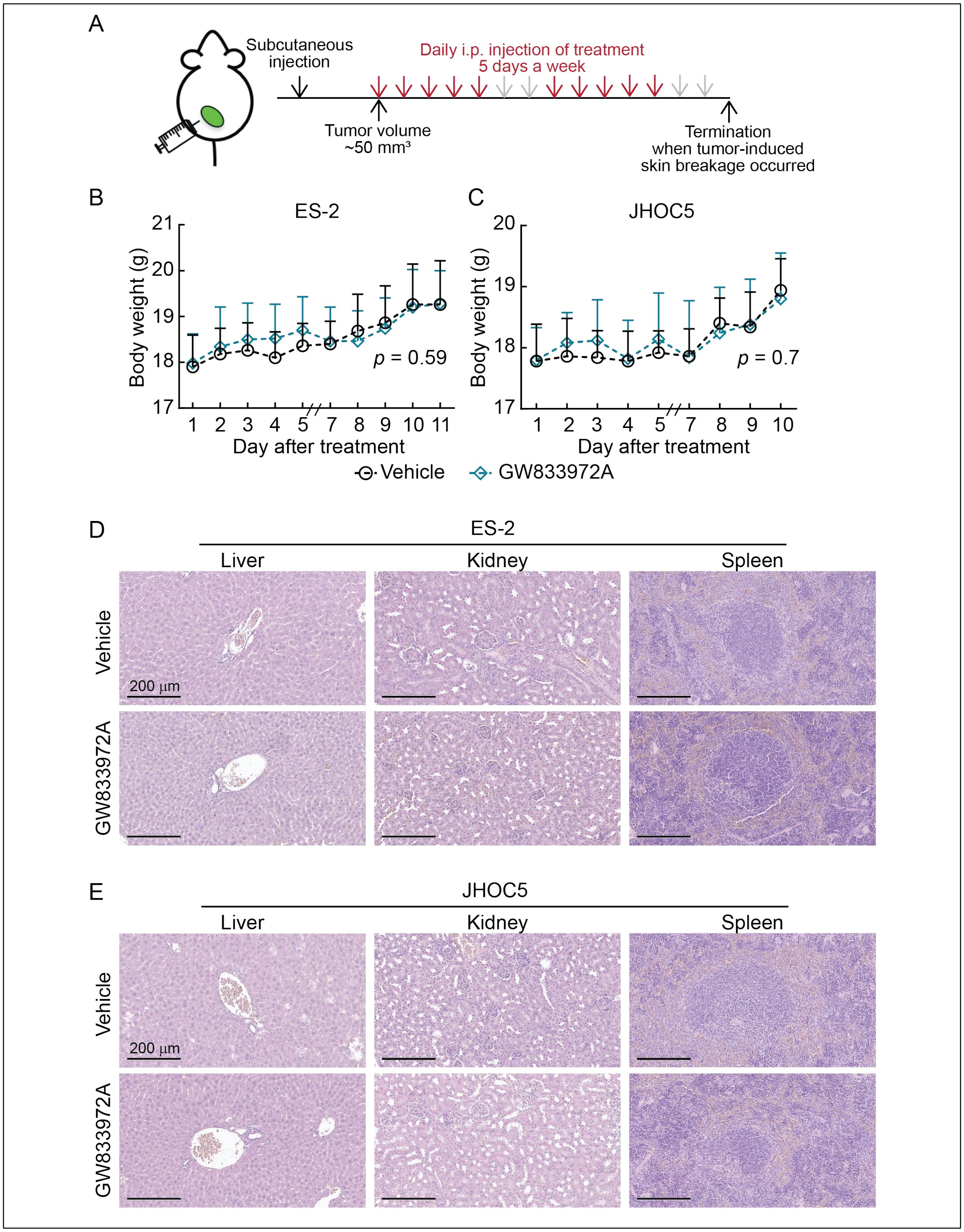

**Figure S7. Subcutaneous xenograft models using ES-2 and JHOC5 cells.**

(A) Diagram of the procedure used for subcutaneous xenografts. $2\times10^6$ ES-2 or $5\times10^6$ JHOC5 cells mixed with Matrigel with 1:1 ratio were subcutaneously injected into the lower flanks of the mouse. When tumors reached 50 $mm^3$, mice were intraperitoneally injected with either vehicle or 10 mg/kg GW833972A daily for five days a week. Mice were euthanized when tumors in the control group reached 2-cm diameter or when the tumor-induced skin breakage occurred in the vehicle control group.

(B) Subcutaneous xenograft model in NUDE mice using ES-2 cells with or without 10 mg/kg GW833972A treatment. Five mice were used for each group. Nonlinear regression (curve fit) analysis was used to test for significant differences between the body wieght curve of the vehicle control and GW833972A treated group (*P*-value is indicated).

(C) Subcutaneous xenograft model in NUDE mice using JHOC5 cells with or without 10 mg/kg GW833972A treatment. Five mice were used for each group. Data formatting is as described for (B).

(D) Representative images of Hematoxylin and Eosin (H&E) staining using liver, kidney and spleen tissue sections from mice bearing ES-2 derived tumors. Scale bars indicate 200 μm.

(E) Representative images of Hematoxylin and Eosin (H&E) staining using liver, kidney and spleen tissue sections from mice bearing JHOC5 derived tumors. Scale bars indicate 200 μm.

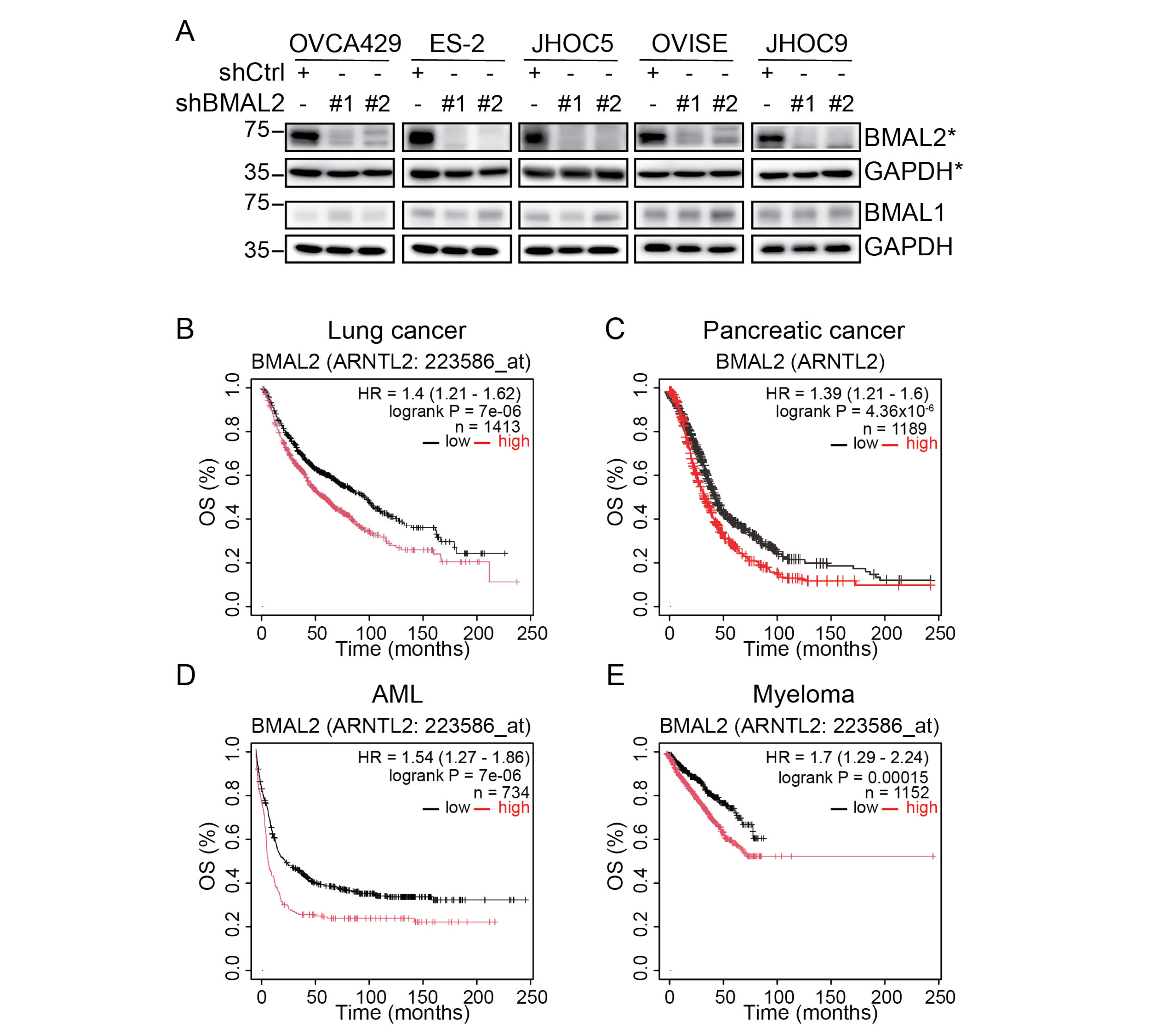

**Figure S8.**

(A) Depletion of BMAML2 did not affect BMAL1 level in OCCC cells. IB of BMAL1 and BMAL2 protein levels with GAPDH as a loading control in OCCC cell lines without (shCtrl ) or with BMAL2 depletion (shBMAL2#1 or #2). Blots shown are from one representative experiment of three replicates. *, BMAL2 blots presented here were from Figure 2B.

(B)-(E) Kaplan-Meier overall survival (OS) analysis of lung cancer (B), pancreatic cancer (C), AML (D) and myeloma (E) cancer patients grouped by BMAL2 expression. The BMAL2 high group is indicated by red line and the BMAL2 low group is indicated by black line. The *P* value was determined by log-rank test. The results shown here are based upon data generated by the KM plotter (https://kmplot.com/analysis/).

### Detailed GW833982A structural characterization

### 1. General Information

##### Nuclear magnetic resonance (NMR) spectra

All reported compounds were characterized by $^{1}H$, $^{13}C$, and $^{19}F$ NMR spectra recorded on a Bruker AVIII HD 400MHz spectrometer. Solvents used to dissolve compounds were indicated in the procedures accordingly. The chemical shifts are reported in ppm and were internally referenced to the residual solvent signals: for $CDCl_3$, δ 7.26 ppm for $^{1}H$ NMR and δ 77.16 ppm for $^{13}C$ NMR; for DMSO-$d_6$, δ 2.50 ppm for $^{1}H$ NMR and δ 39.52 ppm for $^{13}C$ NMR, respectively. The coupling constants (*J*) were reported in Hz, and the splitting patterns were singlet, doublet, triplet, quartet, multiplet, and broad peaks, abbreviated as s, d, t, q, m, and bs, respectively.

##### High-resolution mass spectroscopy (HRMS)

HRMS were obtained on a Bruker microTOF-QII mass spectrometer connected with an Agilent 1260 Infinity HPLC. The ionization source parameters were ESI positive, nebulizer (2.5 Bar), capillary (4.5 kV), dry heater (220 °C), end plate offset (-500 V), scan region (50-3000 m/z), dry gas (8.0 L/min), and collision cell RF (250.0 Vpp).

### 2. Synthesis

##### Materials

All the reagents, solvents, and the starting materials were commercially purchased with high purities and used as received without further purification unless otherwise noted. Benzyl 2-chloro-4-(trifluoromethyl)pyrimidine-5-carboxylate was purchased from Matrix Scientific, 3-chloroaniline, methylene chloride from Thermo Scientific, 4-picolylamine from AK Scientific, ethanol (EtOH) from Shimakyu Chemical, potassium hydroxide from Showa Chemical, and 1,4-dioxane from Fisher Scientific. In general, the progress of the reactions was monitored by thin-layer chromatography (TLC) plates using Merck silica gel 60 $F_{254}$. The handheld UV lamp manufactured by Analytik Jena, UVG-11 (254 nm and 365 nm), was used to identify the spots.

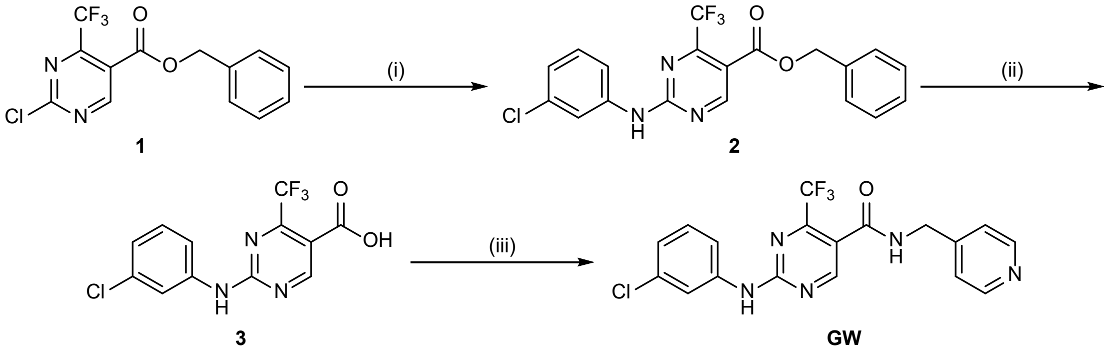

Scheme S1. (i) 3-chloroaniline, 1,4-dioxane, RT, 18 h (ii) KOH, EtOH, reflux, 12 h (iii) EDC·HCl, HOBt·$H_2O$, 4-picolylamine, RT, 18 h.

**Benzyl 2-((3-chlorophenyl)amino)-4-(trifluoromethyl)pyrimidine-5-carboxylate (2)**

A solution of compound **1** (1.27 g, 4.00 mmol, 1.00 equiv.) in 1,4-dioxane (13 mL) was stirred at room temperature under an argon atmosphere, followed by the addition of 3-chloroaniline (2.1 mL, 19.9 mmol, 4.96 equiv.), and the reaction mixture was stirred for 18 h at room temperature. Upon completion of the reaction, 1,4-dioxane was removed under reduced pressure using a rotary evaporator. The resulting solid was redissolved in ethyl acetate (50 mL) and washed with 2 N $HCl_{(aq)}$ (25 mL). The organic layer was dried over anhydrous $Na_2SO_4$, filtered, and concentrated to approximately 5 mL. The resulting mixture was triturated with hexane (170 mL) to afford a white solid, which was collected by suction filtration and dried under reduced pressure to afford a white solid (1.34 g, 3.28 mmol, 82%). $R_f$= 0.36 (EA/Hex = 9/1). $^1$H NMR (400 MHz, $CDCl_3$): $\delta$ 9.06 (s, 1H), 7.81 (s, 1H), 7.77 (bs, 1H, NH), 7.46-7.36 (m, 6H), 7.28 (t, $J$ = 8.1 Hz, 1H), 7.11 (d, $J$ = 7.8 Hz, 1H), 5.38 (s, 2H). $^{13}$C NMR (101 MHz, $CDCl_3$): $\delta$ 162.9, 159.8, 155.6 (q, $J$ = 36.9 Hz), 140.0, 135.0, 134.9, 130.1, 128.8, 128.7, 124.3, 120.1 (q, $J$ = 276.4 Hz), 120.0, 118.0, 114.3, 67.2. $^{19}$F NMR (376 MHz, $CDCl_3$): $\delta$ -66.7. ESI-HRMS calculated for $C_{19}H_{14}ClF_3N_3O_2^+$, [M+H$^+$] 408.0721, found 408.0721.

**2-((3-Chlorophenyl)amino)-4-(trifluoromethyl)pyrimidine-5-carboxylic acid (3)**

To a stirred suspension of compound **2** (1.34 g, 3.28 mmol, 1.00 equiv.) in ethanol (33 mL) was added a solution of KOH (575 mg, 10.2 mmol, 3.12 equiv.) in ethanol (11 mL), and the mixture was refluxed for 12 h under an argon atmosphere using a reflux condenser. After completion of the reaction, the mixture was cooled to room temperature, and ethanol was removed under reduced pressure using a rotary evaporator. The resulting solid was taken up in water (33 mL) and extracted with diethyl ether (50 mL). The combined aqueous layers were then acidified to pH 1 with 12 N $HCl_{(aq)}$, during which a precipitate formed. The solid was then collected by suction filtration and dried under reduced pressure to afford a yellow solid (965 mg, 3.03 mmol, 93%). $R_f$= 0.01 (EA). $^1$H NMR (400 MHz, DMSO-$d_6$): $\delta$ 10.8 (bs, 1H, NH), 9.08 (s, 1H), 7.97 (t, $J$ = 2.0 Hz, 1H), 7.68 (ddd, $J$ = 8.3, 2.0, 0.8 Hz, 1H), 7.37 (t, $J$ = 8.1 Hz, 1H), 7.12 (ddd, $J$ = 8.0, 2.0, 0.8 Hz, 1H). $^{13}$C NMR (101 MHz, DMSO-$d_6$): $\delta$ 164.1, 162.8, 159.6, 153.6 (q, $J$ = 35.5 Hz), 140.4, 133.2, 130.2, 122.8, 121.7, 120.3 (q, $J$ = 276.1 Hz), 118.3, 114.5. $^{19}$F NMR (376 MHz, $CDCl_3$): $\delta$ -65.6.

**2-((3-Chlorophenyl)amino)-*N*-(pyridin-4-ylmethyl)-4-(trifluoromethyl)pyrimidine -5-carboxamide (GW)**

A suspension of compound **3** (965 mg, 3.03 mmol, 1.00 equiv.) in methylene chloride (30 mL) was stirred at room temperature under an argon atmosphere and were added 1-hydroxybenzotriazole hydrate (701 mg, 4.58 mmol, 1.51 equiv.), 1-(3-dimethylamino-propyl)-3-ethylcarbodiimide hydrochloride (883 mg, 4.60 mmol, 1.52 equiv.), *N*,*N*-diisopropylethylamine (1.6 mL, 9.19 mmol, 3.03 equiv.), and 4-picolylamine (0.46 mL, 4.53 mmol, 1.50 equiv.) sequentially, and the resulting reaction mixture was stirred for 18 h at room temperature. Upon completion of the reaction, methylene chloride was removed under reduced pressure using a rotary evaporator. The resulting solid was redissolved in ethyl acetate (70 mL) and extracted sequentially with water (50 mL) and saturated $NaHCO_{3(aq)}$ (50 mL). The combined organic layers were dried over anhydrous $Na_2SO_4$, filtered, and concentrated to approximately 5 mL. The resulting solution was triturated with hexane (150 mL) to afford a white solid, which was collected by suction filtration and dried under reduced pressure to afford a white solid (1.22 g, 2.99 mmol, 98%). $R_f$= 0.33 (EA). $^1$H NMR (400 MHz, DMSO-$d_6$): $\delta$ 10.6 (bs, 1H, NH), 9.23 (t, $J$ = 6.0 Hz, 1H, NH), 8.93 (s, 1H), 8.53 (dd, $J$ = 4.4, 1.6 Hz, 1H), 7.97 (t, $J$ = 2.0 Hz, 1H), 7.66 (dd, $J$ = 8.3, 1.9 Hz, 1H), 7.39-7.34 (m, 3H), 7.10 (ddd, $J$ = 8.0, 2.0, 0.8 Hz, 1H), 4.49 (d, $J$ = 6.0 Hz, 2H). $^{13}$C NMR (101 MHz, DMSO-$d_6$): $\delta$ 164.3, 160.6, 159.4, 152.1 (q, $J$ = 36.9 Hz), 149.9, 148.1, 141.1, 133.5, 130.6, 122.7,

122.5, 120.7 (q, $J$ = 276.9 Hz), 119.7, 119.2, 118.2, 42.2. $^{19}$F NMR (376 MHz, $CDCl_3$): $\delta$ -65.6. ESI-HRMS calculated for $C_{18}H_{14}ClF_3N_5O^+$ [M+H$^+$] 408.0833, found 408.0830.

### 3. NMR spectra

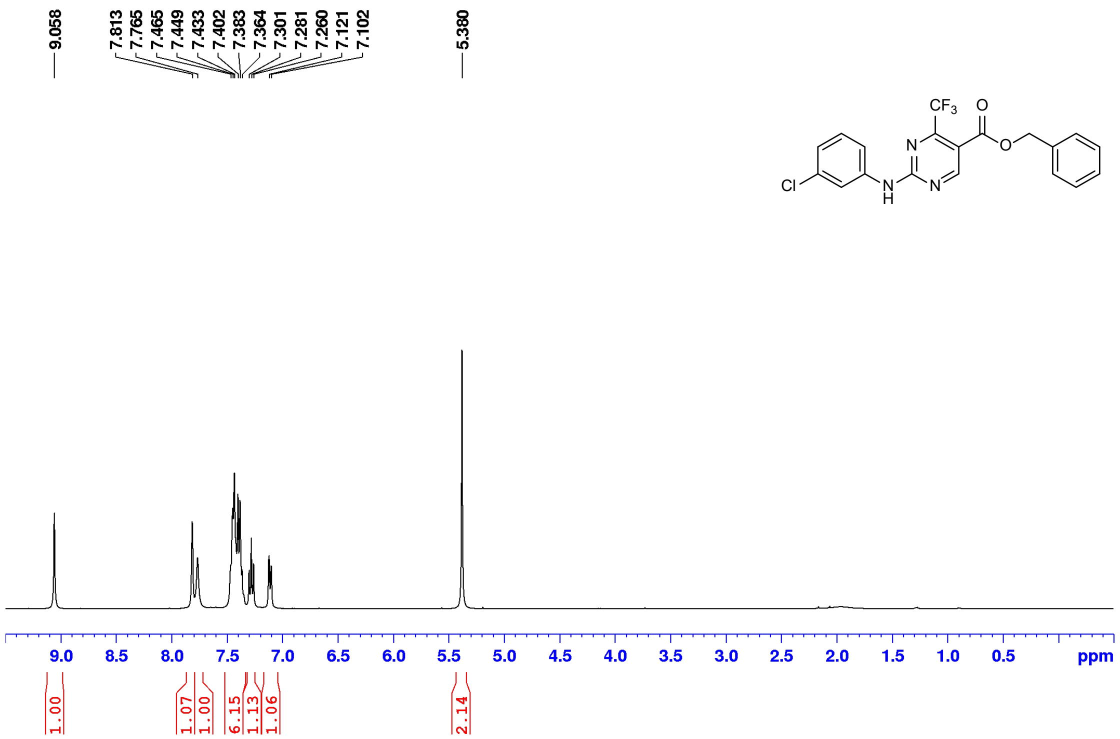

**Figure S9.** $^{1}$H-NMR spectrum of benzyl 2-((3-chlorophenyl)amino)-4-(trifluoromethyl)pyrimidine -5-carboxylate (**2**) in $CDCl_3$.

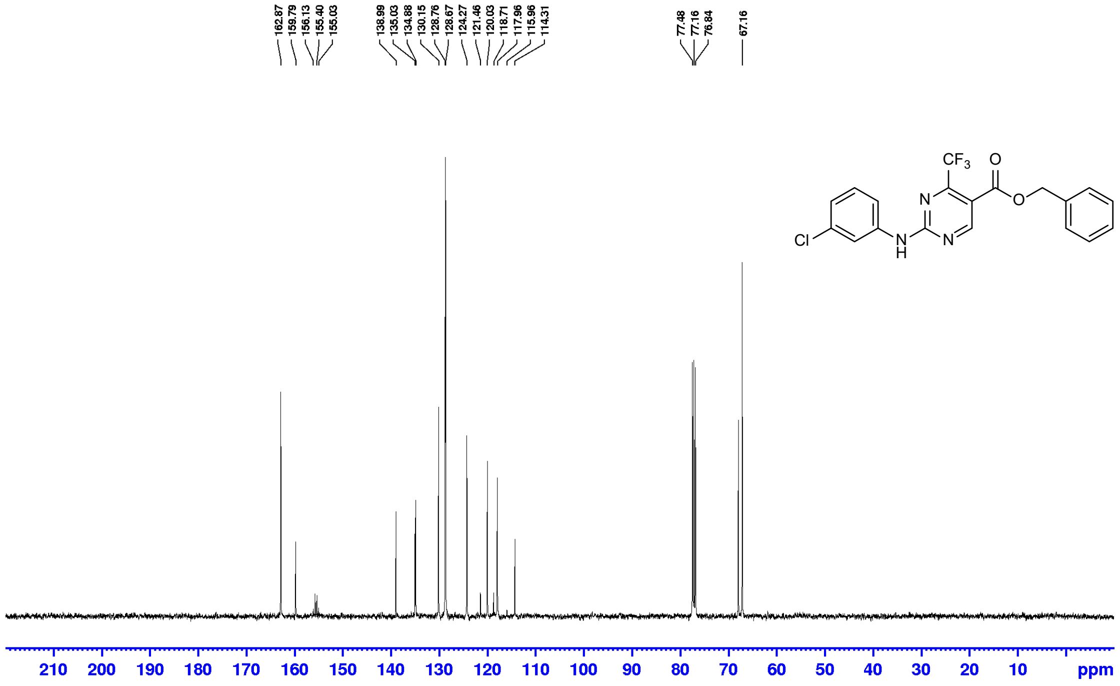

**Figure S10.** $^{13}$C-NMR spectrum of benzyl 2-((3-chlorophenyl)amino)-4-(trifluoromethyl)pyrimidine -5-carboxylate (**2**) in $CDCl_3$.

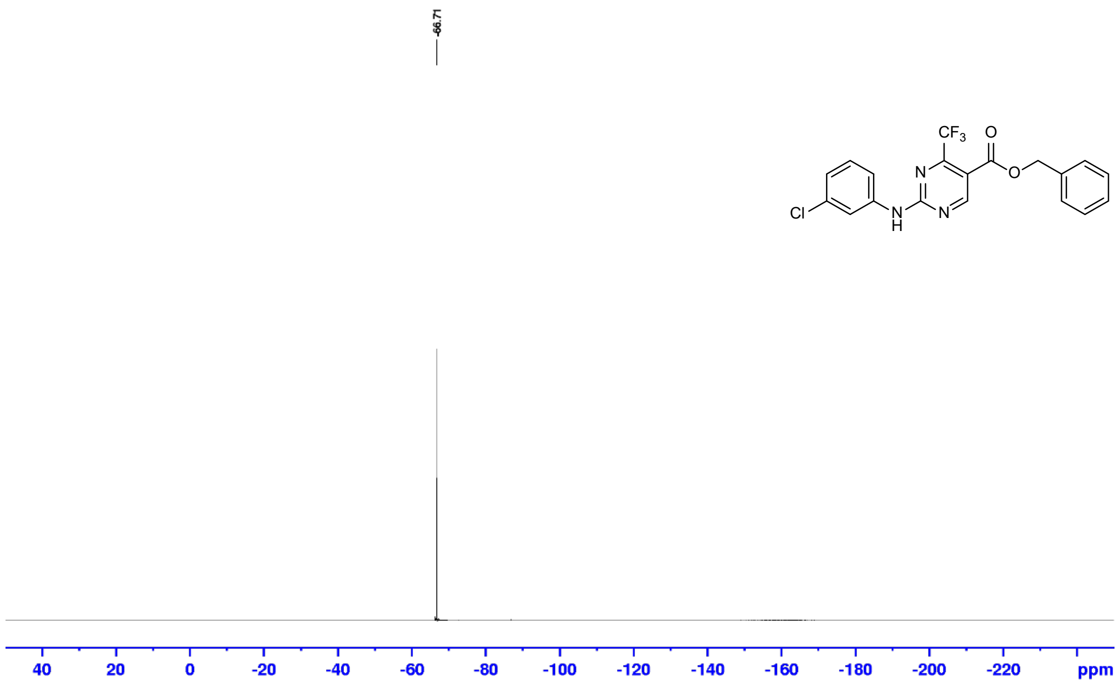

**Figure S11.** [19]F-NMR spectrum of benzyl 2-((3-chlorophenyl)amino)-4-(trifluoromethyl)pyrimidine -5-carboxylate (**2**) in $CDCl_3$.

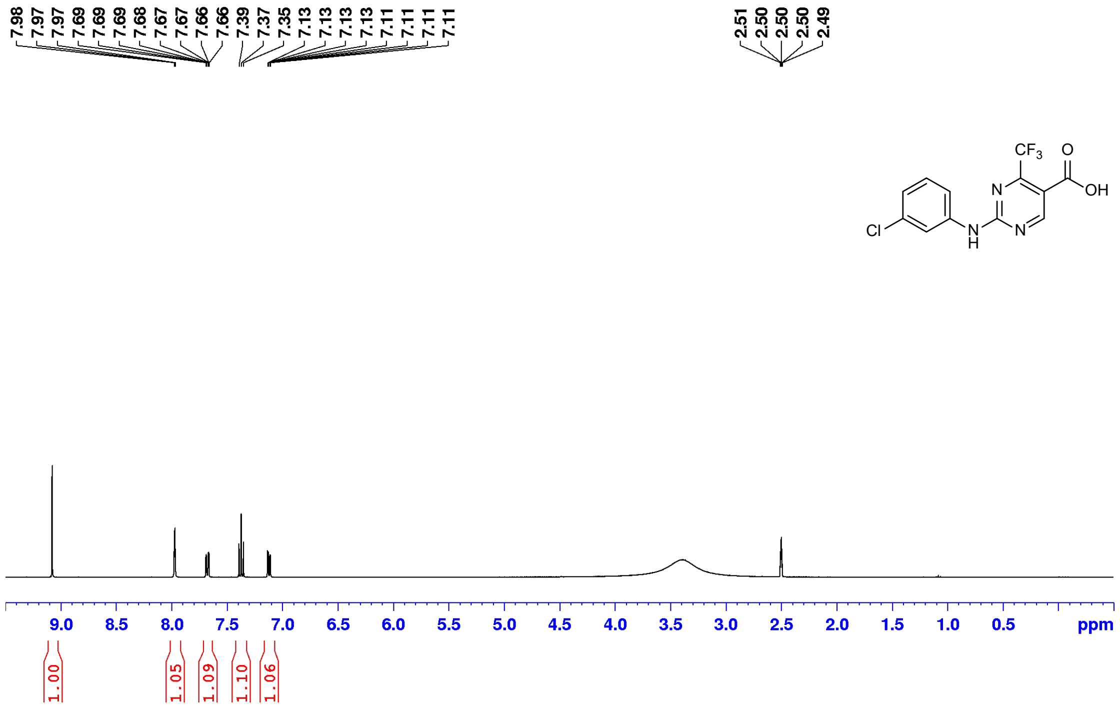

**Figure S12.** [1]H-NMR spectrum of 2-((3-chlorophenyl)amino)-4-(trifluoromethyl)pyrimidine-5-carboxylic acid (**3**) in DMSO-$d_6$.

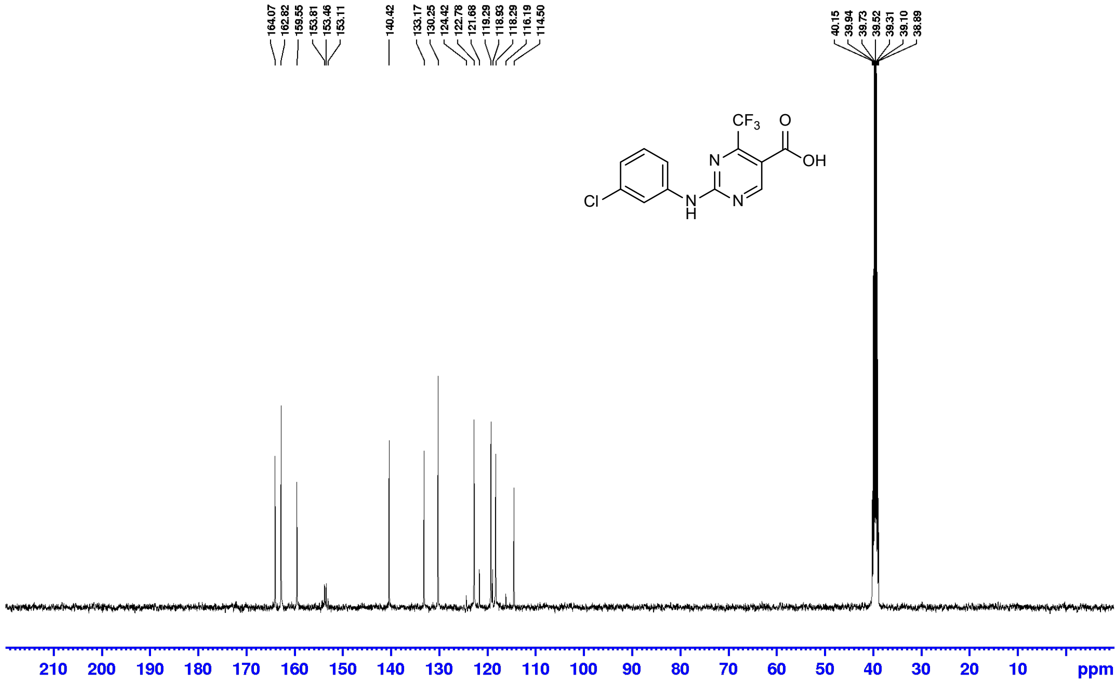

**Figure S13.** $^{13}$C-NMR spectrum of 2-((3-chlorophenyl)amino)-4-(trifluoromethyl)pyrimidine-5-carboxylic acid (**3**) in DMSO-$d_6$.

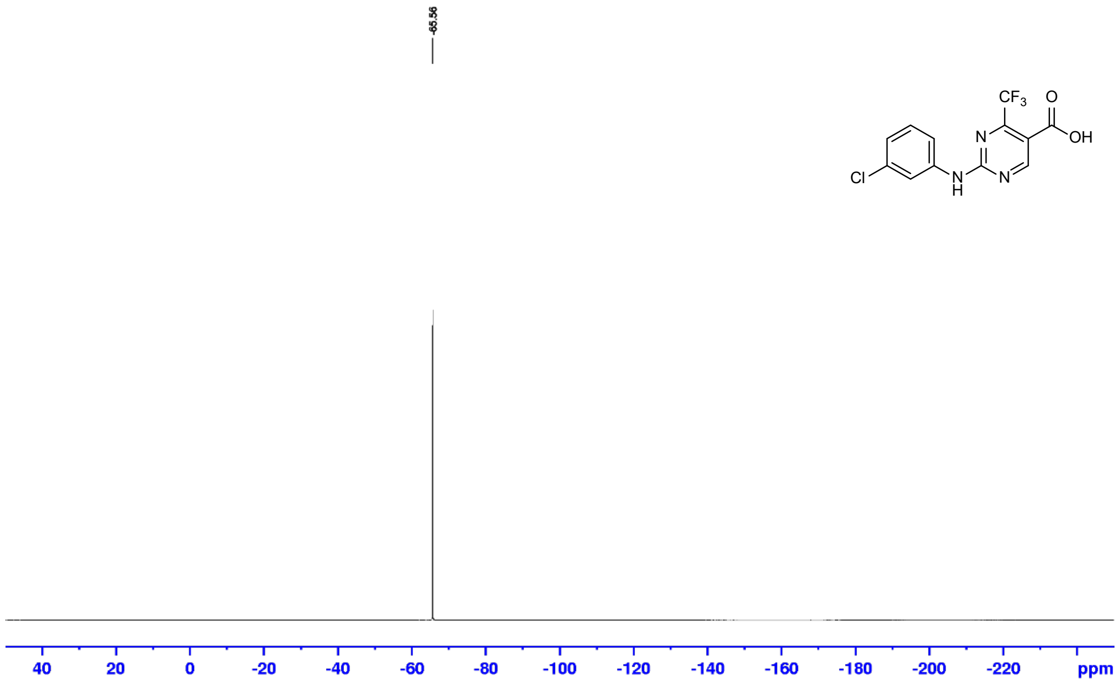

**Figure S14.** $^{19}$F-NMR spectrum of 2-((3-chlorophenyl)amino)-4-(trifluoromethyl)pyrimidine-5-carboxylic acid (**3**) in DMSO-$d_6$.

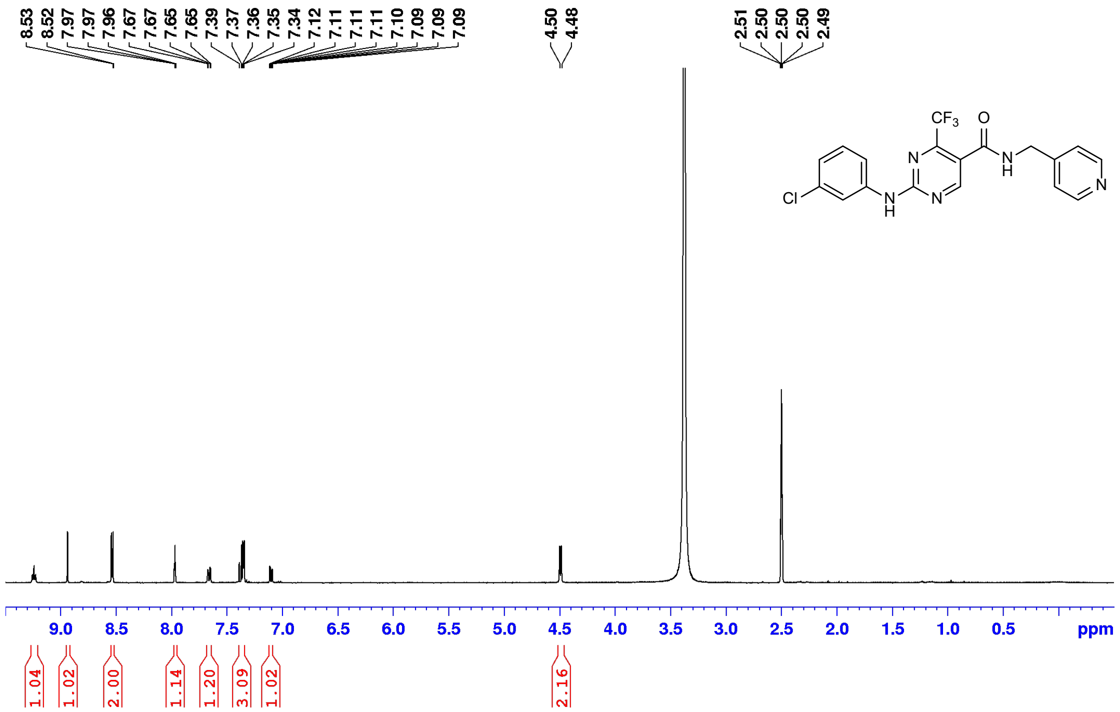

**Figure S15.** $^{1}$H-NMR spectrum of 2-((3-chlorophenyl)amino)-*N*-(pyridin-4-ylmethyl)-4-(trifluoro methyl)pyrimidine -5-carboxamide (**GW**) in DMSO-$d_6$.

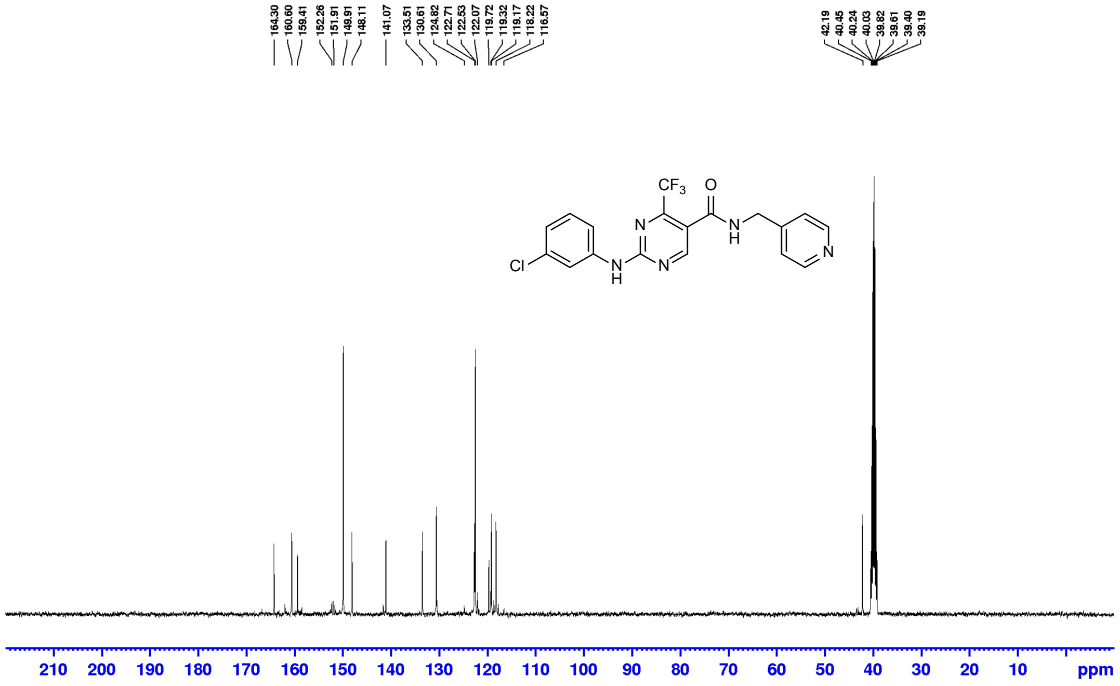

**Figure S16.** $^{13}$C-NMR spectrum of 2-((3-chlorophenyl)amino)-*N*-(pyridin-4-ylmethyl)-4-(trifluoro methyl)pyrimidine -5-carboxamide (**GW**) in DMSO-$d_6$.

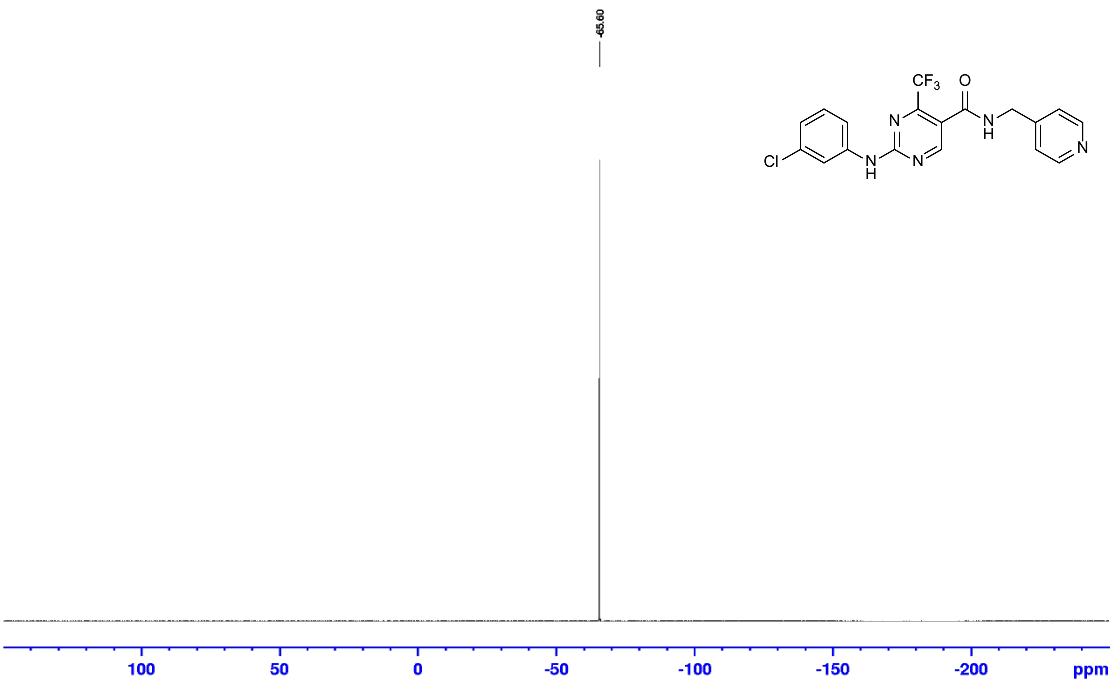

**Figure S17.** [19]F-NMR spectrum of 2-((3-chlorophenyl)amino)-*N*-(pyridin-4-ylmethyl)-4-(trifluoro methyl)pyrimidine -5-carboxamide (**GW**) in DMSO-$d_6$.
